# Gene Regulatory Network Inference reveals *TCF4* as a key a player in neuroblastoma gene expression circuitry

**DOI:** 10.64898/2026.07.07.737136

**Authors:** Catherine Koering, Elodie Vallin, Franck Picard, Sandrine Gonin-Giraud, Olivier Gandrillon

**Affiliations:** Univ Lyon, ENS de Lyon, Univ Claude Bernard, CNRS UMR 5239, INSERM U1210, Laboratory of Biology and Modelling of the Cell, 46 allée d’Italie Site Jacques Monod, Lyon, F-69007, France; Inria, Villeurbanne, 69693, France

## Abstract

Neuroblastoma (NB), a pediatric cancer arising from disrupted sympathetic neuron differentiation, exhibits marked heterogeneity and limited therapeutic options.

To better understand its molecular circuitry dynamics, we applied CardamomOT, a novel Gene Regulatory Network (GRN) inference framework, to single-cell RNA-seq data from patient-derived tumoroids. This approach models gene regulation via piecewise deterministic Markov processes, capturing transcriptional bursting and protein-mediated feedback, overcoming limitations of RNA velocity (e.g., gene independence and lack of biological time).

We identified a continuous chromaffin-to-sympathoblast differentiation trajectory along which we selected 85 dynamically relevant genes enriched in cell cycle and DNA replication functions. Notably, 9 genes overlapped with those driving normal sympathoadrenal differentiation, underscoring tumor-normal tissue similarity.

The inferred 85-genes network reproduced quite well experimental gene expression patterns *in silico*, and allowed to predict protein-level dynamics. Furthermore, it allowed to predict the effect of perturbations (both knock-out and overexpression) of hub genes (e.g., *TCF4* and *PLK1*). We show that those perturbations significantly altered cell fate proportions *in silico*, with *TCF4* KO increasing chromaffin-like cells and reducing proliferative late sympathoblasts.

Predictions regarding *TCF4* were tested using drug inhibition as a proxy for the gene KO. Using the BET inhibitor JQ1 indeed induced profound effect on the transcriptomic identity of our tumoroids. Finally cell fate proportions were also altered *ex vivo* closely resembling the predicted output.

Our work therefore demonstrates that NB tumoroids retain a dynamic, differentiation-like architecture amenable to GRN modeling. Predicted druggable targets offer testable therapeutic avenues, including repurposing BET inhibitors or PLK1 inhibitors, potentially in combination.

## Introduction

Neuroblastoma (NB) is the most common extracranial solid tumor in children under five years of age. These tumors typically arise near the adrenal glands and are characterized by marked inter- and intra-patient heterogeneity ([1]). Clinical outcomes vary dramatically, ranging from spontaneous regression in infants to aggressive, treatment-resistant disease in older children. While low- and intermediate-risk cases achieve >90% five-year survival, high-risk cases remain below 60%, even with recent therapeutic advances ([2]). Treatment options remain limited, with few actionable mutations and very few drugs clinically validated ([3]), although GD2-targeted CAR-T cell therapies offer a promising avenue ([4]).

There is therefore an urgent need to better understand the molecular mechanisms driving NB initiation and maintenance, in order to develop effective therapies specifically targeting high-risk disease. NB is a paradigmatic example of a tumor arising from disruptions in a developmental process - specifically, during the maturation of migratory neural crest cells into sympathetic neurons ([5]; [6]; [7]). This deep entanglement between normal differentiation and oncogenesis is not unique to NB and has been observed across multiple tumor types ([8]). It is thus compelling to view NB development and maintenance as emerging from an underlying dynamical system that mimics and distorts developmental differentiation.

Among studies interrogating NB at single-cell resolution (([9]; [10]; [11]; [12]; [13]; [14]; [15]), only a few have explored potential dynamical behaviors. To move beyond static snapshots of differentiation, some authors have applied RNA velocity analysis to infer directional gene expression trajectories ([9]; [11]; [12]). RNA velocity relies on a kinetic model that distinguishes unspliced from spliced mRNA counts and fits gene-specific parameters using an ODE-based transcription model, enabling prediction of future mRNA levels ([16], [17], [18]). However, despite its time-dependent formulation, RNA velocity lacks explicit biological time interpretation ([19]), and models genes independently, ignoring regulatory interactions ([17]).

To overcome these limitations, we resorted to CardamomOT, a novel Gene Regulatory Network (GRN) inference framework that enables a fully mechanistic and dynamical view of gene regulation ([20]; [21]; [22]). In this model, the expression dynamics of each gene is modeled with a stochastic process that captures transcriptional bursting and subsequent protein synthesis ([23]; [24]). Genes are coupled via an interaction function that models how the proteomic environment feeds back into gene burst frequency. Integrating CardamomOT with the simulation engine HARISSA allowed us to infer executable GRN models capable of reproducing observed experimental data ([21]). CardamomOT generates a fully executable model, that can be directly run by a computer system to perform simulations or predictions ([25]).

In the present work, we applied this approach to single-cell RNA-seq (scRNA-seq) data from Patient-Derived Tumoroids (PDTs). PDTs are three-dimensional *ex vivo* structures that recapitulate key histological features of NB. Those have become valuable tools for both fundamental and translational research in NB ([26]; [27]; [28]).

Using CardamomOT, we identified an 85-gene network driving chromaffin-to-sympathoblast differentiation that accurately reproduces expression patterns and predicts the impact of gene perturbations on cell fate. Experimental validation using the BET inhibitor JQ1 confirmed these predictions, significantly altering tumoroid transcriptomes and shifting cell fate proportions *ex vivo*.

Our work therefore demonstrates that NB tumoroids retain a dynamic, differentiation-like architecture amenable to GRN modeling, allowing testable prediction toward potential therapeutic avenues, including repurposing known inhibitors, potentially in combination.

## Results

To study the differentiation dynamics of NB cells we generated scRNA-seq data from Patient-Derived Tumoroids (PDTs). Cells from one patient were obtained as OPDX, injected into mice, generating PDX that were subsequently grown in 3D cultures to generate PDTs. Single-cell RNAseq was performed on those PDTs using RevGel-seq ([29]; see material and methods for a complete description). We generated two batches from the same culture of tumoroids derived from one patient (SJNBL013762) at two different times in culture. The first batch displayed the expression of 60,617 genes in 1744 cells, the second the expression of 60,617 genes in 2601 cells. Since both datasets were clearly not separated by PCA (Supplementary figure 1), nor by UMAP (not shown) we decided to aggregate them into a unique dataset. After filtering (see material and methods) we obtained a matrix that displays the expression profiles of 30,034 genes in 4345 cells.

Most studies apply a clustering algorithm, like Leiden of Louvain, on the raw dataset. Those will give rise to clusters, even in homogeneous cell population. We therefore decided to first assess whether or not there was sufficient structured cell-to-cell variability in our dataset in order to decide if it contained meaningful subpopulations that should be further resolved. For this we applied Phiclust ([30]) which returned a high clusterability index of 0.9992. A value close to 1 means that multiple phenotypic subpopulations are present, so we decided to investigate the presence of relevant subpopulations further. For this we applied the Leiden clustering algorithm (Figure 1A), generating 8 clusters. It was obvious that, except for cluster 6 (see below), all clusters were connected through a continuous variation. The next question was then whether or not those clusters should be considered as resulting from a single dynamical process generating the cell diversity within our dataset. This would be in line with the hypothesis that neuroblastoma arise from the subversion of a differentiation process (see discussion for details). We therefore first applied the RNA velocity approach using scVelo ([31]) to compute a pseudotime representing each cell’s position along a continuous differentiation axis (Figure 1B). This allowed to define a left to right differentiation sequence in the UMAP space. We then used CellRank ([32]) to confirm the existence of a clear differentiation direction (Figure 1C). Trajectories dynamics was further confirmed using scFates ([33]) combining CellRank and a simple principal tree algorithm (Figure 1D).

**Figure 1:**
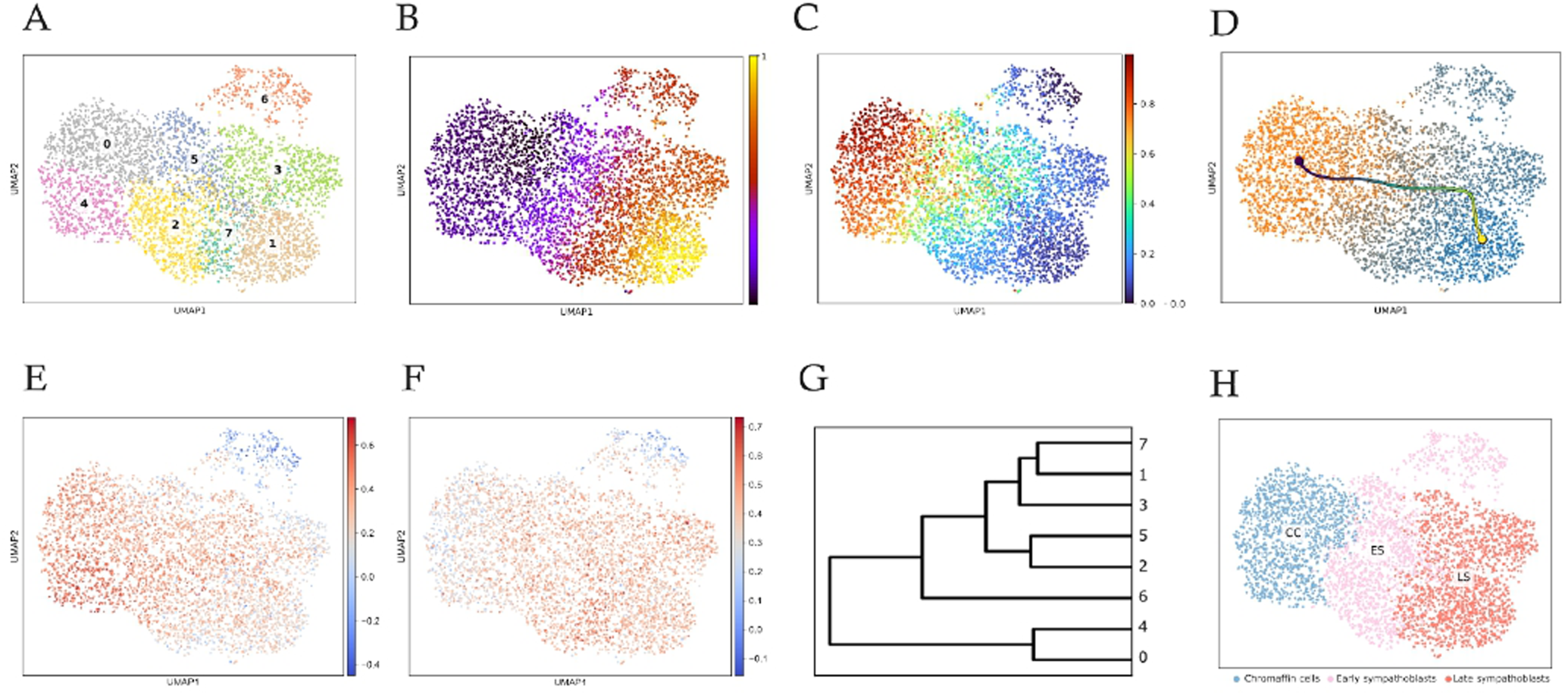
UMAP representation of PDTs single-cell RNAseq. A: The dots were color-coded according to their clustering identity assessed by the Leiden algorithm ([34]) with a resolution of 0.8 using 2000 Highly Variable Genes. B: Each cell is color-coded according to its defined velocity pseudotime as computed by scVelo ([31]; early pseudotimes are labelled in black, late in yellow). C: Each cell is color-coded according to its probability to be at the beginning of the underlying dynamical process, as assessed by CellRank ([32]; high probability is labelled in red). D: The left-right trajectory was constructed using scFates, starting in cluster 0 and ending in cluster 1. E: Each cell is color-coded according to the Kameneva’s chromaffin cells gene expression signature. F: Each cell is color-coded according to the Kameneva’s sympathoblasts gene expression signature. G: Hierarchical clustering of the 8 Leiden clusters. H: The final cluster-based definition of our three cell types.

The next question consisted in identifying relevant cell types from this clusters. There is no established consensus as to what cell types can be found in neuroblastoma in the literature (see discussion for details). We therefore first applied signature genes specific for normal cell types as defined in ([9]; see material and methods for the list of genes). We observed cells expressing chromaffin-specific genes at the initiation of the differentiation process, whereas the end of the sequence was characterized by cells expressing sympathoblasts-specific genes (Figure 1E and 1F). Interestingly, this was also observed using two other gene signatures (Supplementary figure 2). Such an uninterrupted continuum of transcriptional cell states connecting chromaffin cells to sympathetic phenotypes was previously observed in mice TH-MYC cells ([12]).

We then performed hierarchical clustering from the Leiden clusters (Figure 1G) to see if some clusters could be regrouped. Clusters 0 and 4 were close in the hierarchical tree, and harbored the strongest chromaffin signature. We therefore decided to regroup them under the “chromaffin cells” label. At the right extremity of the UMAP, clusters 3, 7 and 1 were close in the hierarchical tree, and harbored the strongest sympathoblast signature. We therefore decided to regroup them under the “Late sympathoblast” label. The remaining intermediary clusters 6, 5 and 2 were pooled as early sympathoblasts (see below and Discussion section for this choice of cell types).

In order to explore the robustness of that result to the clustering algorithm, we compared Leiden’s clustering to Louvain clustering (Supplementary figure 3). Overall, the cell type definition was quite stable with a pretty robust definition of chromaffin cells, and a more variable definition of early versus late sympathoblasts, as expected from the proximity of those two cell types in the hierarchical tree. The impact of cell type definition on the output of the GRN simulation has been explored below (see Supplementary figure 4).

Of note, cluster 6 stands somewhat outside of the main differentiation trajectory. Thorough investigation demonstrated that this cluster was composed in part from cells harboring the sympathoblasts signature and cells harboring a Schwann cell precursors-type signature (Supplementary figure 5 and discussion for its role in the process).

We then characterized the molecular composition of the 3 cell types we just defined. The chromaffin type was characterized by (i) an early velocity pseudotime (Figure 2A), a low cell cycle index (Figure 2B), a specific gene expression pattern (2C-G), including a low *MYCN* expression. This is in line with chromaffin cells being less proliferative that sympathoblasts ([35]).

**Figure 2:**
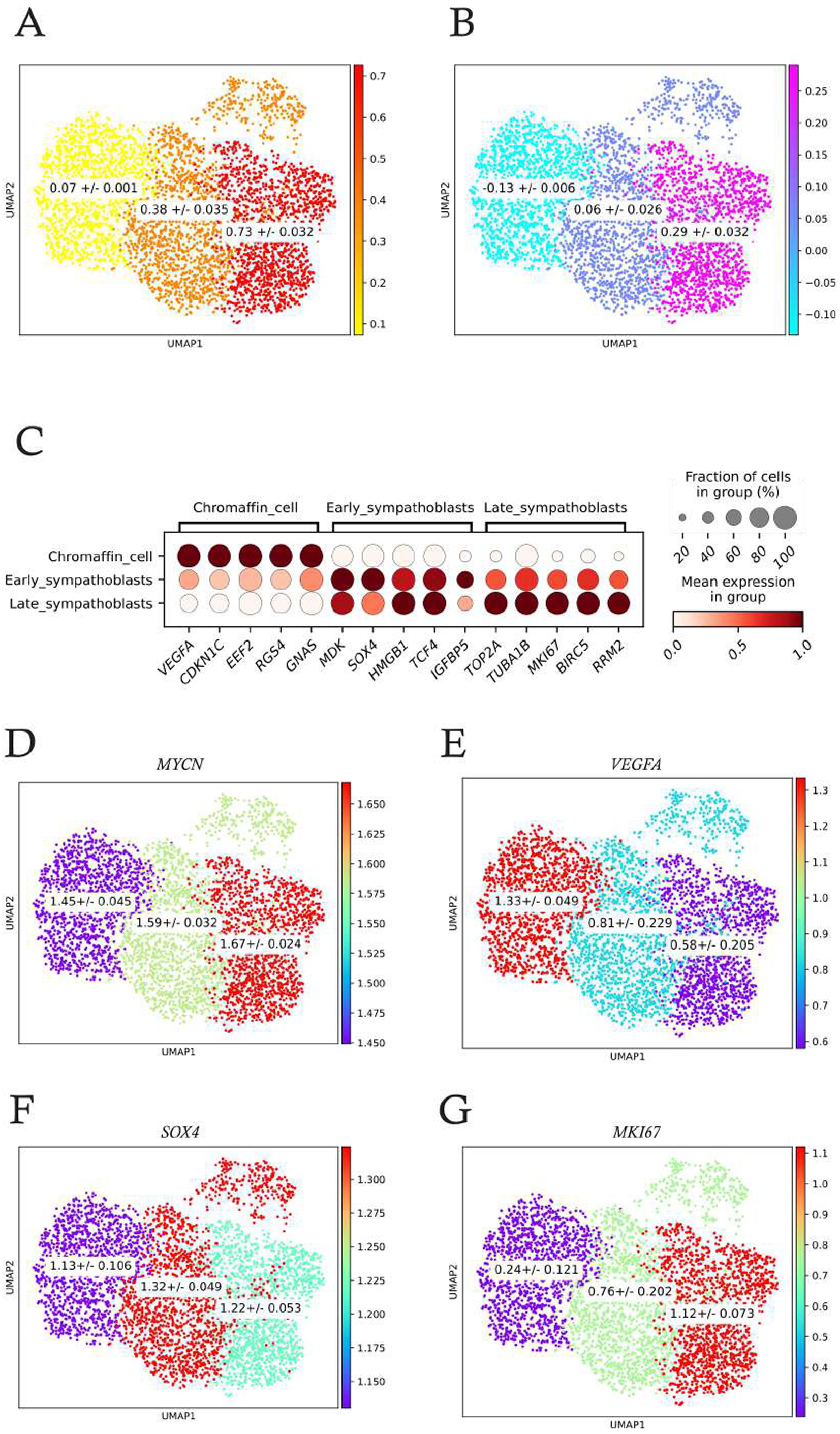
Molecular characterization of the three cell types. A: Mean velocity pseudo-time per cell type. B: Mean S_score per cell type. C: Differential gene expression using the rank_genes_groups function from scanpy ([36]). D to G: Mean expression of *MYCN* (D); *VEGFA* (E), *SOX4* (F) and *MKI67* (G) per cell type. Expression values have been log1p transformed and they range between 0 and 2.18. A mean expression of 1 means that the gene is moderately expressed. In order to see a cell-by-cell variability in gene expression, see Supplementary figure 6.

Altogether our analyses are coherent with a continuous dynamical process stemming from non-proliferating *VEGFA*^+^ chromaffin cells and proceeding toward *MYCN* ^+^ proliferating sympathoblasts.

The application of CardamomOT requires time-stamped single-cell transcriptomic data (see Disucssion section). We therefore defined 5 « times » of evolution of the underlying dynamical process ranging from the lowest to the highest velocity pseudotime values (Figure 3A), and fixed the value of one time slice to 12.5 hours (see Discussion below).

**Figure 3:**
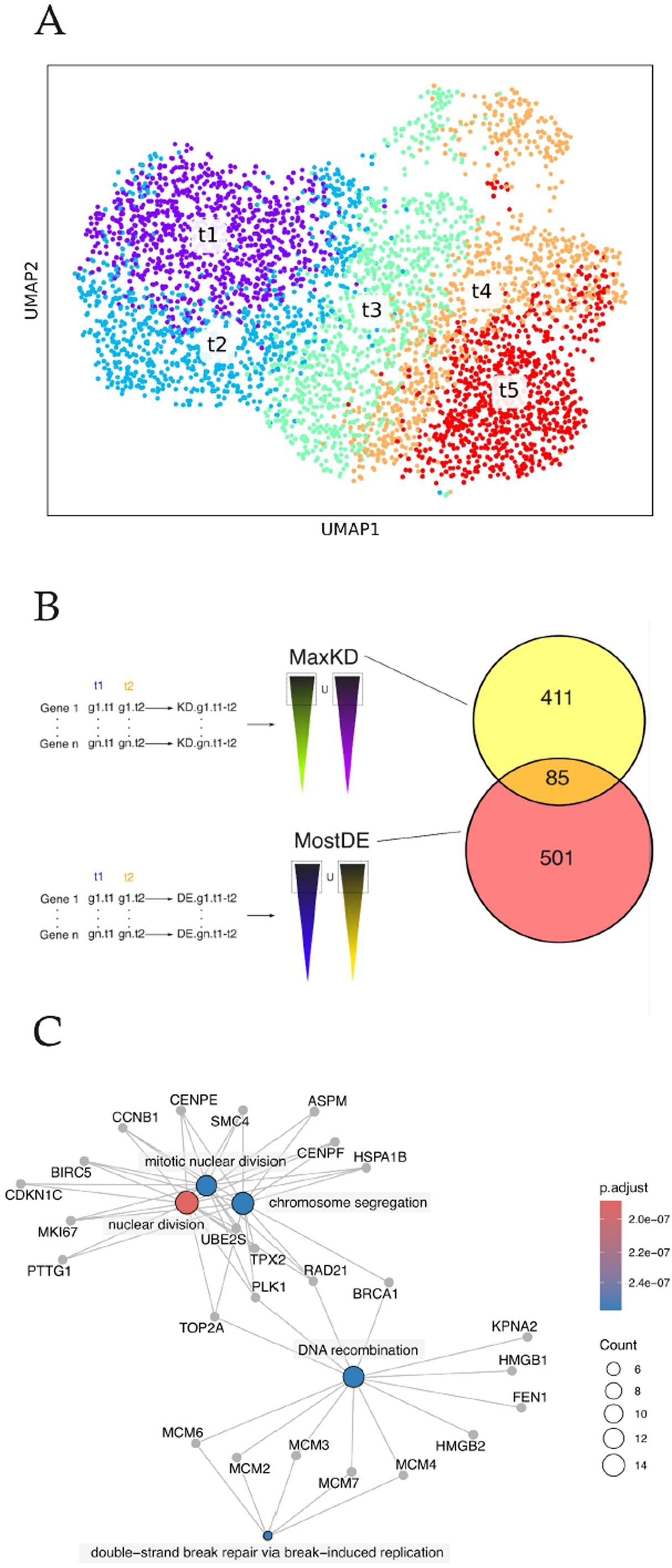
A: UMAP positioning of the 5 times defined along the velocity pseudo-time sequence. B: The gene selection procedure. We first computed a Kantorovitch distance for all genes between 2 consecutive time points. We then took the union of the 200 genes harboring the largest distance for each delta-time, which left us with a list of 586 genes (some genes might be among the most distant between two or more time points), labelled MaxKD. Very similarly, we computed an entropy value for all genes between 2 consecutive time points. We then took the union of the 200 genes harboring the largest entropy for each delta-time, which left us with a list of 496 genes labelled MostDE. The intersection of those two lists gave us a list of 85 genes (Supplementary Table 1) that were used for inferring the GRN. C: Gene Ontology (GO) analysis of the 85 gene list (see supplementary file at https://osf.io/gkh89/ for quantitative details).

We then explored the most relevant gene set to be used for the GRN inference (Figure 3B). Because our inference and simulation scheme is fundamentally rooted in exploiting the power of distributions ([21]; [37]), we first computed a distance between gene expression distribution using Kantorovich distance ([38]) for all genes at all delta-times.

Similarly, we have shown that another relevant metrics was delta-entropy. This computes the difference in entropy of the distribution of a gene between two time points, revealing how much the shape of the distribution has changed. We have shown that the genes displaying the highest difference, called the most delta-entropic genes, were shown to represent the most relevant genes for the differentiation process ([39]). We therefore also computed delta-entropy for all genes for all delta-times. The intersection of those two lists provided us with a list of 85 genes that was used for inferring the GRN (Supplementary table 1). This list of 85 genes contains *MYCN* and *PHOX2B*, two genes known to be important for NB development ([40]; [41]; [42]; [43]; [7]; [44]). Similarly, both *DLK1* ([45]) and *MEG3* ([46]), two genes involved in cancer stemness, were present in our list.

Six Long Non Coding RNAs (lncRNAs; *GAS5, SNHG7, ZFAS1, MEG3, H19, MIAT*) were among the selected genes. Although the presence of noncoding genes among genes simulated using a two states model with protein production might see counterintuitive (see Discussion below), we nevertheless kept those genes in the light of their known importance in various cancers ([47]; [48]). More specifically, among the 6 lncRNAs, both *MEG3* ([49]) and *MIAT* ([50]) were shown to be involved in neuroblastomas.

We analyzed potential functional overrepresentation among those 85 genes using clusterProfiler ([51]). Three out of five of the most overrepresented GO biological processes pertained to cell division (Figure 3C). DNA recombination including 5 members of the MCM (minichromosome maintenance) family was also significantly overrepresented. The most overrepresented GO molecular function pertained to single-stranded DNA helicase activity, once again driven by a large presence of members of the MCM family (see supplementary file at https://osf.io/gkh89/).

We finally assessed the overlap between this list made from tumor cells with the one we previously published to infer the normal sympathoadrenal differentiation in human ([52]). 9 genes were common between those two lists (*CHGA, DLK1, TH, HMGB2, IGFBP2, MDK, MEG3, RGS4* and *VIM*) highlighting the close proximity between the tumor cells and their normal counterpart.

We then used CardamomOT to infer a GRN ([20]). Results are displayed in Figure 4A. The parameter *θ*_j,*i*_ (see material and methods) defines the strength of the interaction going from gene j to gene i (outgoing for j and incoming for i). We could then compute the number of interactions per gene. We observed the expected (see e.g. [52] or [20]) striking difference between the distribution of the numbers of outgoing and incoming nodes (Figure 4B). The number of incoming nodes was distributed according to a normal (i.e., Gaussian) distribution, whereas the outgoing nodes displayed a very heavy long tail toward higher values, as well as a large number of low or null values. Such a distribution suggests the existence of hub genes and is characteristic of small-world networks, a disputed notion in biology [53]. The intensity of all outgoing interactions for the 14 most densely connected genes is shown in Figure 4C. Most genes had either only positive or negative interactions, with few, like *DNAJB1*, harboring both types.

**Figure 4:**
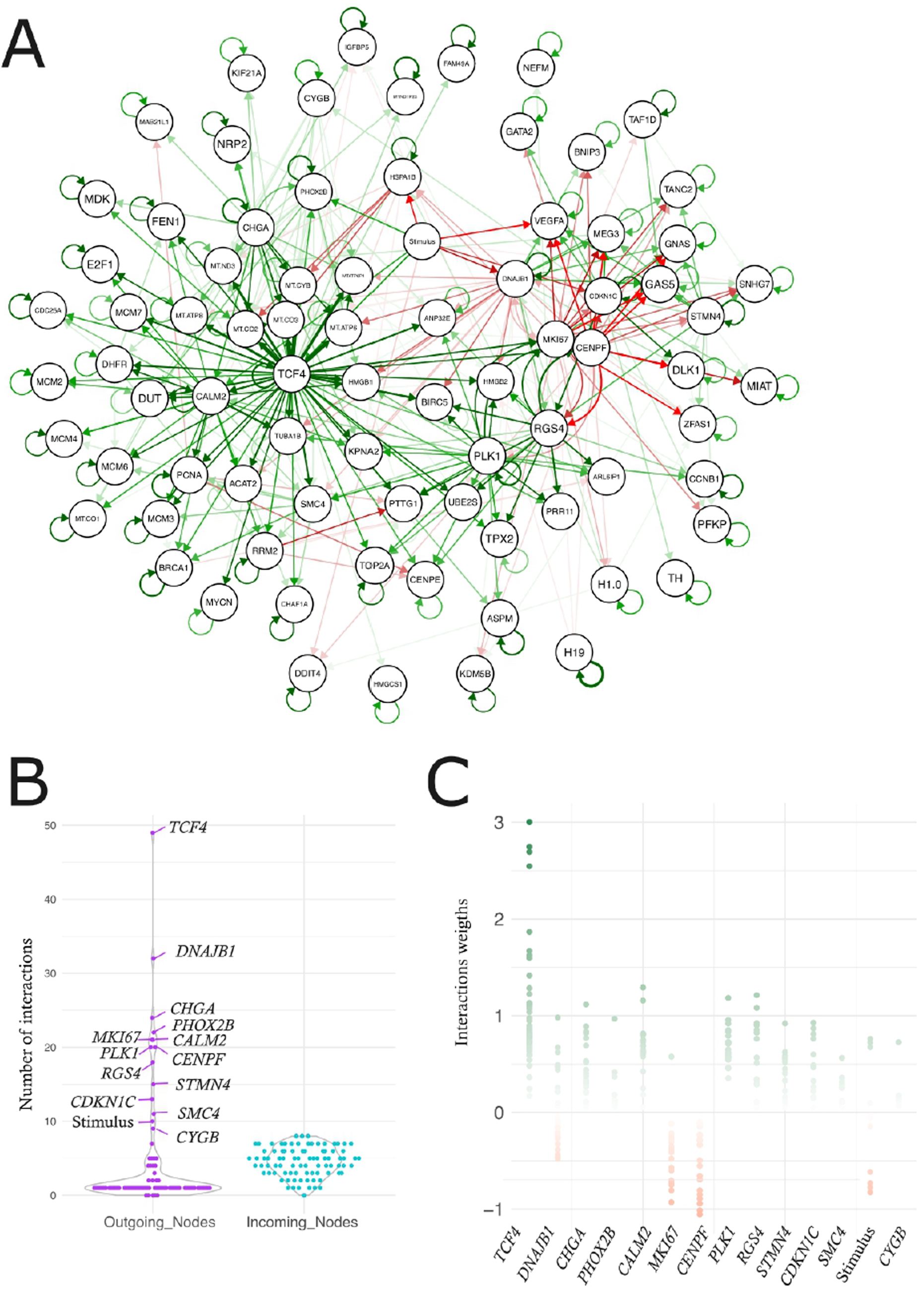
A: The inferred GRN. Positive regulations are displayed in green, negative one in red. B: Violin plot representing the number of outgoing or incoming interactions per gene. C: The actual values of the outgoing interactions are shown for the 14 genes (including the stimulus) with the highest number of interactions.

We then explored the GRN for the presence of specific dynamical motifs, which is a small set of recurring regulation patterns with each motif carrying out specific information-processing functions ([54]).

The first type of motif we analyzed was toggle switches, where two genes inhibit each other. For this we identified pairs of genes i,j such that *θ_i_*_,j_ < 0 and *θ*_j,*i*_ < 0. We identified 2 linked toggle switches connecting *CENPF, DNAJB1* and *MKI67* (Figure 5A). This double toggle switch induced a very clear pattern in the dataset with a respective exclusion of cells expressing *DNAJB1* (in the chromaffin cells) and cells expressing *CENPF and MKI67* (in the late sympathoblast cells).

**Figure 5:**
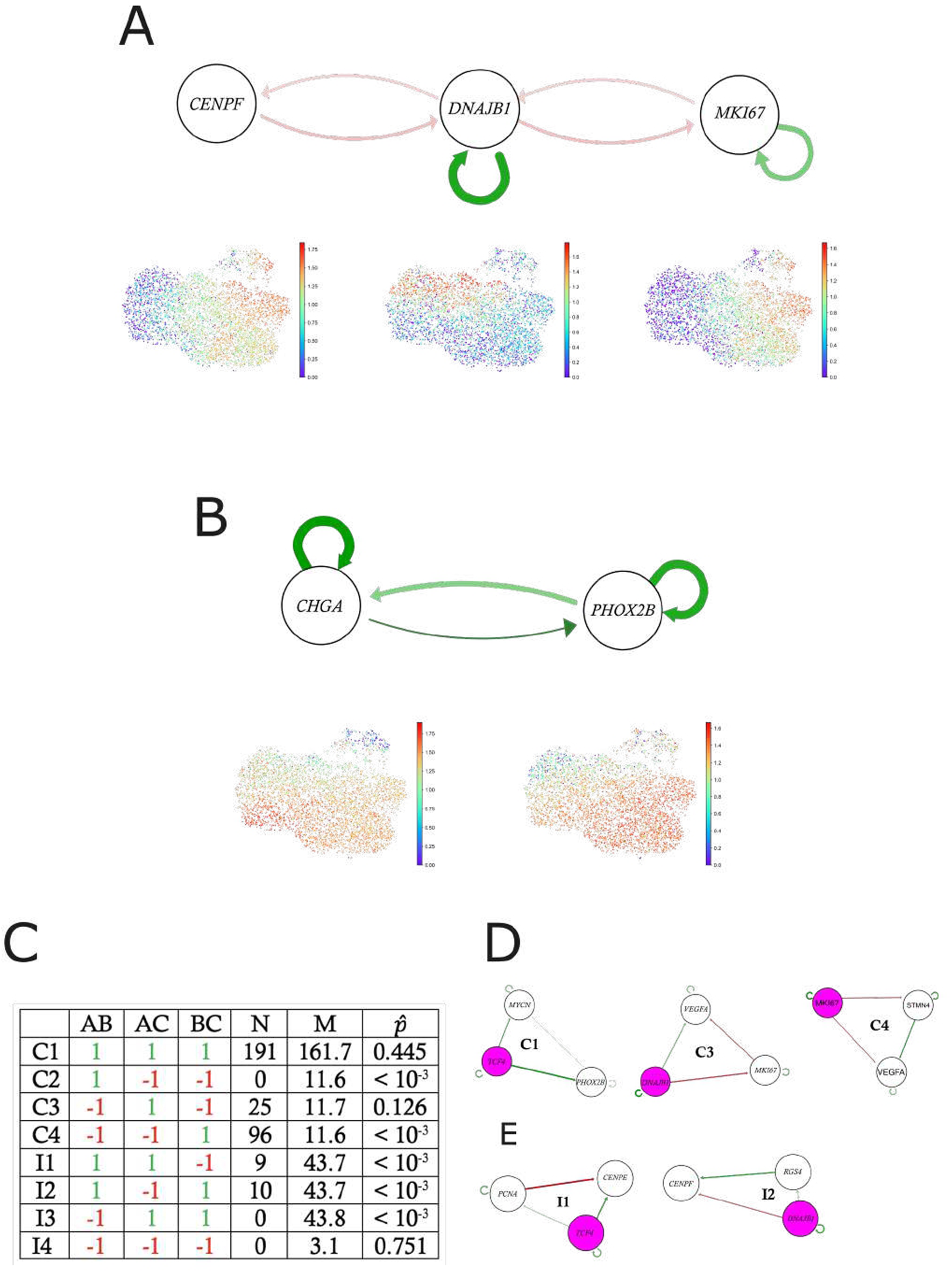
Dynamical motifs. A: A double toggle switch connecting *CENPF, DNAJB1* and *MKI67*.

We also found the existence of one positive reenforcing loop, connecting *CHGA* and *PHOX2B* (Figure 5B), the effect of which could be seen in their co-expression pattern on the UMAP.

We then investigated the existence of feed-forward loops (FFL). Those are 3-node regulatory motif where a regulator affects a target through two paths with identical final effect, therefore leading to an enforced effect. The effects of such motifs are expected to be more subtle than cross-positive or cross-negative loops and are thought to be important for the dynamical behavior of the GRN.

We observed the presence of FFL of the C1, C3 and C4 sort, with the notable exclusion of any C2 type (Figure 5C). We computed an empirical pvalue to assess the significance of the over- or under-representation of each motif. C4 motifs were extremely enriched, whereas C2 motifs were significantly underrepresented.

We finally explored the existence of incoherent feed-forward loops (IFFL). Those are 3-node regulatory motif where a regulator affects a target through two paths with opposing signs, leading to non-monotonic or transient dynamics in the target’s response ([55]). Only two types of motifs out of four (I1 and I2) were present. I1, I2, and I3 were found to be significantly underrepresented (Figure 5C), which was not the case for the I4 motif.

In conclusion, our GRN exhibits a highly structured architecture where simple negative (C4) feed-forward loops serve as the fundamental building blocks, while certain complex configurations of negative loops (C2) are actively excluded. Most but not all of the IFFLs are also actively excluded.

Whether this highly structured architecture results from biological constraints or are at least partly due to some bias in the inference process will require more analysis. The fact that at least one IFFL (type I4) and two FFLs are not significantly different from random, pleads for a *bona fide* biological function.

The observation that C4 (negative FFL) is highly enriched might suggests the network is optimized for temporal delays or noise filtering during differentiation ([56]; [54]).

The respective gene expression pattern is displayed on the UMAP below. B: A positive reenforcing loop connecting *CHGA* and *PHOX2B*. The respective gene expression pattern is displayed on the UMAP below. C: Table displaying the 8 types of loops studied: 4 feed-forward loops (FFL, C1 to C4) and 4 incoherent feed-forward loops (IFFL, I1 to I4). N: number of motifs detected in our GRN. M: mean number of motifs obtained by 10^5^ permutations. *p̂*: estimated pvalue. D: One example of each of the 3 FFLs detected. E: One example of each of the 2 IFFLs detected. The A node is colored in pink. Contrary to Figure 4A, the strengths of the interactions are not drawn at scale for visualization purposes.

One of the main advantages of CardamomOT is that it produces executable GRN models, the time-dependent evolution of which can be simulated, and the resulting simulated dataset can then be compared with the experimentally observed dataset.

In order to proceed with the simulation, one needs to provide CardamomOT with an empirical estimate of the characteristic time process. Once an appropriate timescale has been selected (hours to days), we fined tuned these estimates through the optimization of the fit quality (see below). This led us with a 5 time points sequence: 0, 12.5, 25, 37.5 and 50 hours.

To assess the quality of the GRN-based simulation, we compared the UMAPs between experimental and simulated data (compare Figure 6A and 6B), and we also computed the distances of mRNA distributions (Figure 6C). Both criteria show the close proximity between distributions. The repartition of cell identities also confirmed the quality of the simulation (compare Figure 6D and 6E), although our modeling scheme tended to overestimate the amount of early sympathoblasts.

**Figure 6:**
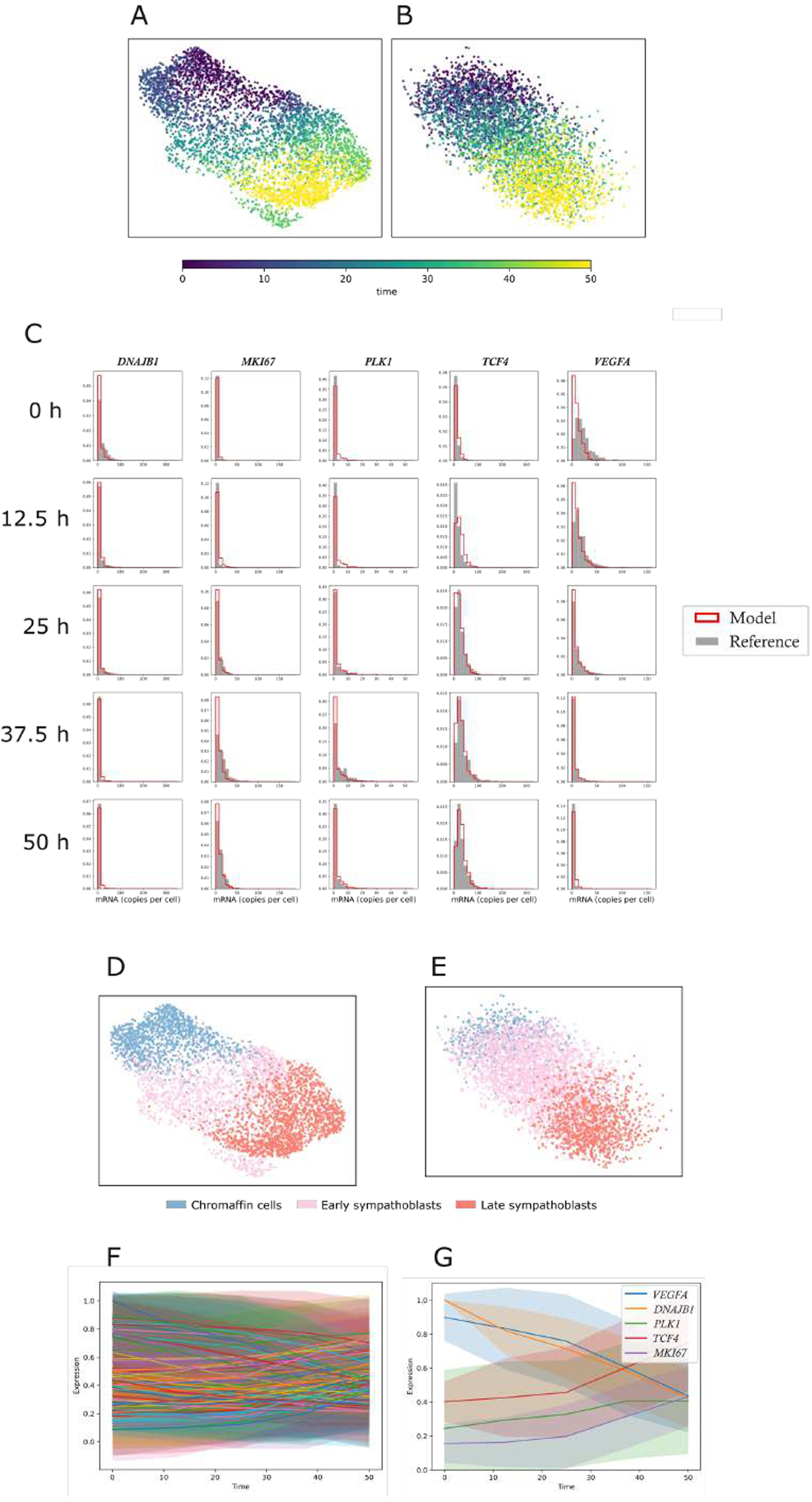
UMAP representation of the dataset used for GRN inference (A) and of the dataset obtained through GRN simulation (B). Cells are color coded according to the model time, in hours. C: Comparison between the experimentally observed mRNA distribution and the simulated distribution from a few selected genes. The marginals computed by the model are shown by red bold-delimited bars, the experimentally-observed marginals are shown as plain bars in grey. A full distribution comparison for the 85 genes can be found at: https://osf.io/gkh89/. D and E: The cells were color-coded through their cell identity, either defined on the original dataset (D) or by random forest classification of the cells produced by the simulation (E). F and G: Time-dependent evolution of the mean+/-SD of all proteins expression (F) or of a selected subset (G). The mean is shown as a plain line, the SD as a dimmed-colored envelope.

Since our model is built upon a gene expression framework in which protein synthesis is explicitly represented, one can also gain access to its behavior at the protein scale. A careful examination of the average behavior of all 85 proteins revealed two distinct classes of genes: those whose expression increased and those whose expression decreased over the modelling time (see Figure 6F for all proteins and 6G for a selected subset). This can be seen also as a validation of the time chosen (cf above) since shorter times (say minutes) would not allow any variation in the amount of proteins to be seen.

Altogether the results of the simulation demonstrate the ability of CardamomOT to correctly reproduce the experimental data, as well as allowing insight into variables that are not experimentally accessible like the distribution of single-cell amount of proteins. The fit quality of the experimental dataset was sufficient to engage in testing the impact of targeted gene alterations on the global behavior of our GRN.

We performed a perturbation analysis of the 15 genes with the highest number of outgoing interactions (Figure 4B), which showed that multiple perturbations could induce a significant variation in the cell-type composition (Figure 7A). We could see a general trend between the most out-connected genes and the most influential ones, as expected. Interestingly, the poorly connected nodes of the network like *CYGB* or *H1.0*, have no influence when perturbed. It is interesting to note that the protein of some of these genes are known to be druggable ([57], see Discussion).

**Figure 7:**
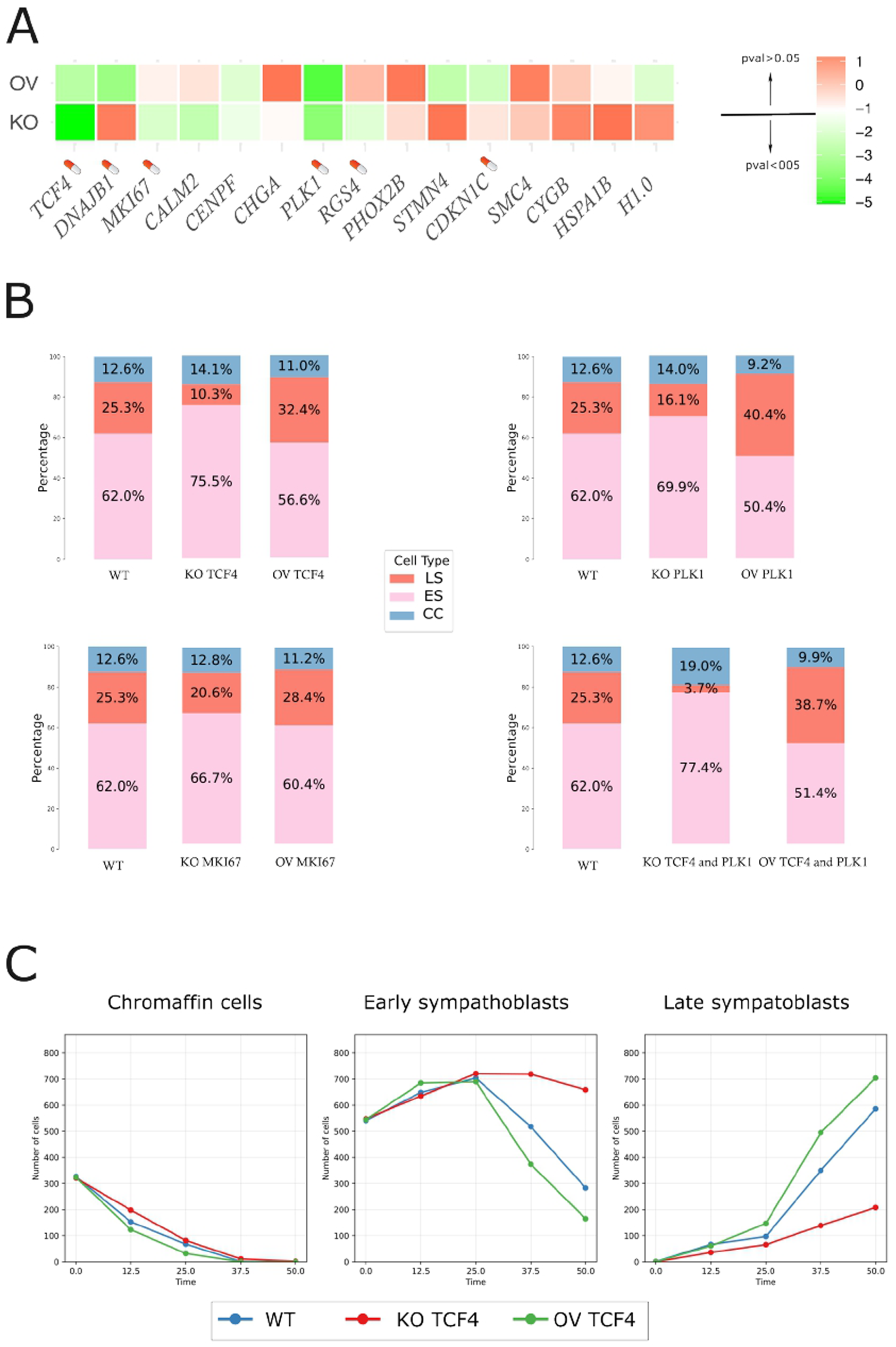
*In silico* perturbations. A: Color coded result of a chi-square test to ascertain the significance in the difference in cell type composition between the WT and the perturbed situation. Bonferroni correction for multiple testing was applied. For representation purposes, the adjusted pvalue was compressed using a double-log scale, and the significance threshold was fixed to 0.05. Genes labeled with a drug capsule are those whose gene products are known to be druggable ([57]). B: Cell type repartition either in the WT simulation or during a single or a double perturbation. CC: chromaffin cells; ES: early sympathoblasts; LS: late sympathoblasts. C: Time-dependent evolution in the amount of cells in the three cell types categories, in the WT case or during *TCF4* perturbations.

In Figure 7B is shown the side-by-side cell type repartition *in silico* between a WT and a perturbed simulation. The KO of *TCF4* induced a strong increase in the amount of early sympathoblasts cells, at the expense of late sympathoblasts. The overexpression of that gene induced a symmetrical effect with a strong increase in the amount of late sympathoblasts. A similar effect was noted for the *PLK1* or the *MKI67* genes although slightly attenuated (Figure 7B). Interestingly, a double perturbation affecting both *TCF4* and *PLK1* produced an additive effect.

Since our framework is based on a dynamical system, we can see how the different cell types evolve as a function of time. Figure 7C shows that for the WT situation, one start with a mix of chromaffin and early sympathoblasts. The process ends with a full late sympathoblasts population, through an intermediary transitory population of early sympathoblasts.

The effect of the simulated *TCF4* KO consists in slowing down the process resulting in a large amount of early sympathoblasts at the latest time. The *TCF4* overexpression had a symmetrical effect, i.e. tends to accelerate the process, resulting in the earlier disappearance of chromaffin cells.

We then explored the robustness of our analysis scheme. For that we defined three ways of slicing the velocity pseudotime: in five equal times, as above, in three equal times or in ten equal times. The gene list retained for GRN inference being computed by variation between time points (see Figure 3B above), we first investigated the overlap between those lists. We observed both a significant overlap as well as some differences (Supplementary figure 4A). Interestingly the TCF4 gene was retained in all three lists. We then simulated the output of the three GRNs and computed their ability to correctly reproduce the cell types, either defined on the basis of the Leiden clustering or on the basis of the Louvain clustering. A quite correct ability to fit was observed in all cases (Supplementary figure 4B). Altogether these results show the robustness of our analysis scheme, with some associated variations.

The next step would consist in experimentally verifying some of the model predictions, as we performed in ([58]). For example, one could knock-out *TCF4* in our tumoroids and verify that we indeed observe the predicted reduction in the amount of proliferating late sympathoblasts. Such a drive of sympathoblasts toward a chromaffin-like state could represent a promising therapy ([12]).

We used specific drugs as a shorter-term alternative, since *TCF4* gene-regulating activity can be targeted indirectly by using bromodomain and extra-terminal domain inhibitors (BETis), like JQ1, known to disrupt TCF4-dependent super-enhancers ([59]). Furthermore, JQ1 has been shown to inhibit human neuroblastoma cell lines growth and induces their differentiation ([60]).

We first analyzed the effect of JQ1 in cell numbers and we observed a reduction that became significant at 48 hours (Figure 8A). This strong effect on cell numbers could either result from a simple antiproliferative effect, or resort to a differentiation blockade effect, as proposed by our modelling approach. In the second case, one could predict that JQ1 would affect both proliferation-related but also non-proliferation-related genes. Cell numbers alteration would be a resulting effect of favoring slowly proliferating immature cells. The timing observed in cell number reduction would argue in favor of the second hypothesis.

**Figure 8:**
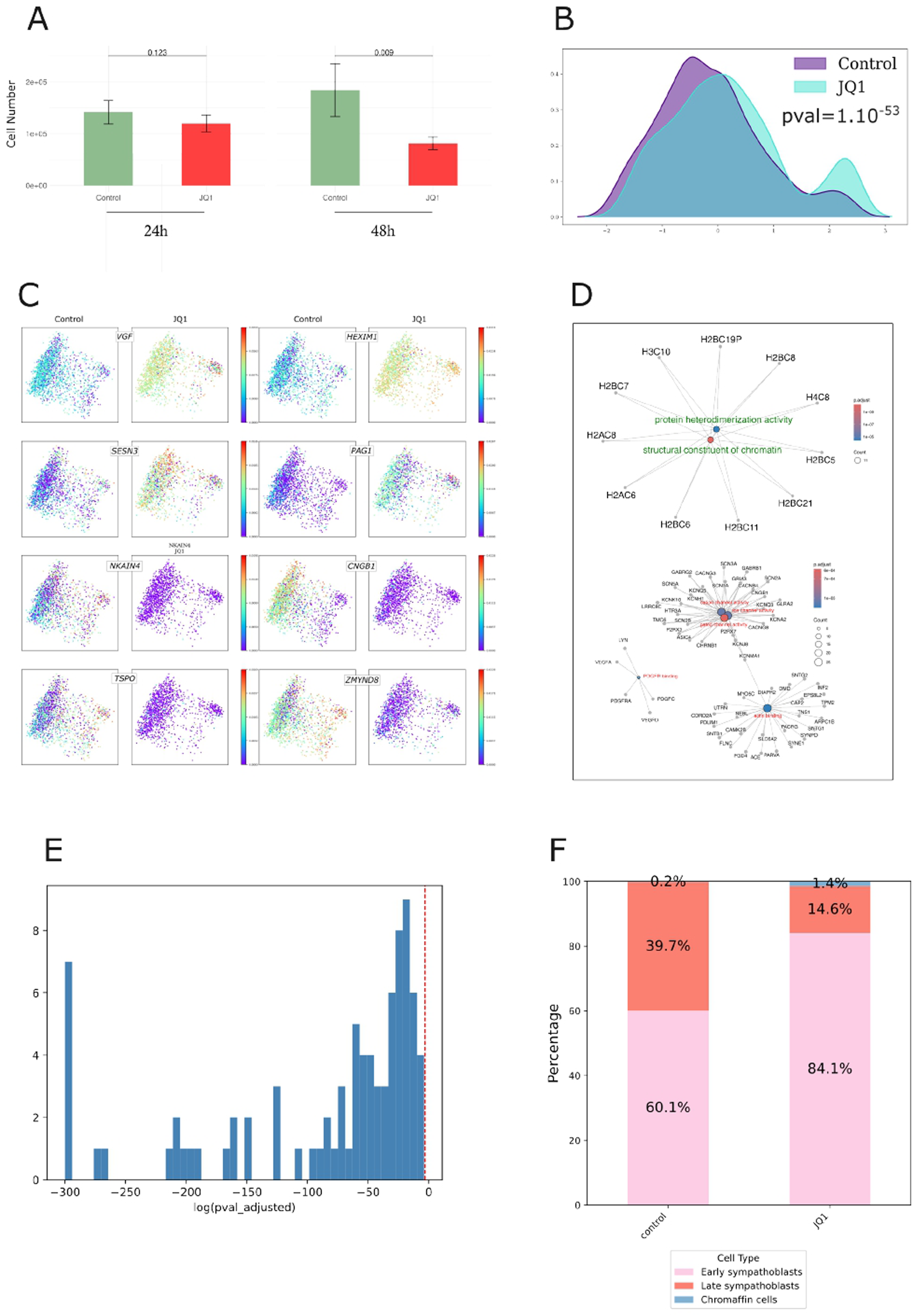
*Ex vivo* testing of *in silico*-predicted perturbations. A: Number of cells per PDTs either untreated or treated with 10^-6^ M JQ1 and counted at either 24 or 48 hours. n=5. pvalue was assessed using a student’s t test; B: Output of the ktest analysis. Cells are represented along the most discriminant axis found by the algorithm; C: Cells are represented along a 2D decomposition of the most discriminant axis found by the algorithm and color coded according to their expression level of genes either up- or down-regulated under the influence of JQ1; D: GO molecular function overrepresentation of up- (in green) or down-regulated (in red) genes. E: Differential expression of the 85 genes of the GRN assessed using ktest. For each gene, we tested H0 being: “there is no significant difference in the expression level of that gene between control and JQ1 treated cells”. Some pvalues being null, we plot the logarithm of the pvalue adjusted for multiple testing plus a small non null value of 10^-300^; F: Cell type composition of PDTs either untreated or treated with JQ1.

To distinguish between those two possibilities, we first compared the full transcriptome of control and JQ1-treated cells using ktest, a kernel-based approach for single-cell transcriptome comparison ([61]; Figure 8B). We observed that JQ1 treatment induced a highly significant change in the overall gene expression pattern in our PDTs (p-value=4.10^-51^). ktest can also be applied on a gene-by-gene basis. We selected genes with an adjusted p-value of less than 10^-3^ and a log foldchange of more that 1, resulting in 120 JQ1-induced genes and 484 JQ1-repressed genes. Plotting the expression value of some of those genes in the kernel-embedding space (Figure 8C) illustrates the amplitude of the variation. We finally assessed a possible GO overrepresentation among those genes (Figure 8D).

Among the JQ1-induced genes, two major molecular functions were overrepresented: protein heterodimerization activity and structural constituent of chromatin. Interestingly all of the genes involved code for histone proteins, involved in both functions. This is in line with the known epigenetic action of JQ1, developed as an epigenetic reader inhibitor ([62]).

Among the JQ1-repressed genes, three major functions emerged: channel activity, actin binding and PDGF-R binding. No strong cell proliferation signature was noted (Figure 8D and see https://osf.io/gkh89/ for an extensive description of GO results) therefore favoring a large impact of JQ1 on cell states beyond simply reducing their proliferation rate. This was further confirmed by the fact that all 85 genes composing our GRN, including but not limited to the predicted TCF4 target genes, were affected (Figure 8E).

We finally assessed the effect of JQ1 treatment on the cell type composition (Figure 8F). We first were confronted with a difficulty: applying the random-forest-learned cell type expression pattern to the control dataset yielded somewhat different quantitative composition, with very few chromaffin cells, illustrating the difficulty in generating standardized samples, even using *ex vivo* approaches. Nevertheless, when we compared the JQ1 and the control samples, we observed the predicted significant decrease in the amount of late sympathoblasts, mostly in favor of early sympathoblasts and a modest but noticeable increase in the amount of chromaffin cells, qualitatively fitting our model’s predictions. The difference between prediction and experimental testing could be partly due to the fact that drug-induced inhibition will not lead to a complete absence of protein activity, in contrast with a knock-out. This nevertheless confirmed our major prediction that JQ1 should have a significant impact on PDTs behavior beyond simply affecting their growth.

## Discussion

In the present work, our main findings are as follows:

Our first contribution is to show that PDTs preserve a high degree of cellular diversity and therefore constitute a powerful framework to investigate differentiation processes and their alteration in NB. This is evidenced, for example, by the high clusterability index reported by Phiclust, and further supported by the presence of multiple cell types - ranging from chromaffin cells to sympathoblasts, and from non-proliferating to highly proliferating cells.

Our second key conclusion is that this variability can be organized along a defined chromaffin-to-sympathoblast axis, revealing the underlying dynamic process.

Third, under the assumption of this dynamic process, we were able to infer a specific GRN that accurately reproduces the time-dependent behavior observed in experimental data.

Fourth, using *in silico* predictions, we could identify genes whose perturbation would most significantly affect the overall dynamical behavior of the tumoroids.

Fifth, we verified experimentally most of CardamomOT-based prediction regarding a key role for the *TCF4* transcription factor regarding cell type composition in our PDTs.

Our analysis fully aligns with the view that NB is the result of the subversion of a normal developmental process, where cell states transitions occur only among states related in the developmental map but not necessarily in the same direction (see below).

Our NB dataset analysis successfully identified the three cell types crucial to sympatho-adrenergic differentiation: chromaffin cells, sympathoblasts, and Schwann cell precursors (SCPs). This is in contrast with a recent study ([63]) where NB presented with a pure committed chromaffin-cell-like phenotype. In this study, authors identified differentiated chromaffin cells (DCHCs). Among the signature genes of DCHCs, two (*NTRK1* and *PCP4*) were clearly not expressed in our NB dataset, highlighting basal differences between the two studies. Whether or not this reflects the natural variation of NB composition among patients will await further investigations.

This is also somewhat at stake with the traditional view of the three cell types composition of NB, that is noradrenergic (NOR) N-type, mesenchymal (MES) S-type, and intermediate I-type, first described on NB cells lines ([64]). Whether those cells can interconvert ([65]; [13]) or not ([66]) adds to the difficulty in NB cell type definition. In our case, 90% of our cells coexpressed *PHOX2B* (a NOR marker; [67]) and *VIM* (a MES marker; [65]). The existence of such cells harboring both NOR and MES features has been previously described in human tumors ([13]). This heterogeneity in NB cell type ([44]) echoes to the very high developmental diversity in embryonic neuroendocrine development ([68]).

Although gene selection was performed differently on the normal dataset made from embryonic human adrenal gland development (where cells were organized along a “stemness axis”; [52]) and on the NB dataset (where cells were organized according to their velocity pseudotime), 9 genes (that is about 10%) were common between those two lists: *CHGA, DLK1, TH, HMGB2, IGFBP2, MDK, MEG3, RGS4* and *VIM.* There are nevertheless differences. A clear branching pattern, from SCPs toward chromaffin cells or sympathoblasts was evidenced in the normal dataset ([52]). The situation is quite different in our PDTs, where the main trajectory displayed a progression from non-proliferating *VEGFA*+ chromaffin cells toward *MYCN* + proliferating sympathoblasts, in line with the transition from immature chromaffin cells into sympathoblasts has been documented in the TH-MYC murine model of NB ([12]). In both cases, one observes a very progressive and continuous process, in perfect accordance with the view that “single-cell omics have reframed NB from a binary classification into a developmental continuum of plastic states.” ([69]).

One should note that there are important discrepancies in the literature regarding the normal differentiation sequence ([9], [11], [10], [7], [5]; [15]; [68]) making the direct comparison with our results quite challenging. It is clear that the generation of neuroblastoma cells cannot be resumed as a simple differentiation blockade in an immature SCP state (this work and [10]). However, studies have confirmed the direct differentiation of immature chromaffin cells into sympathoblasts during human adrenal development ([9]; [11]), a process also observed in the TH-MYC murine model ([12]). In this last case, the authors state that “the recurrent observations of a low-proliferative chromaffin phenotype connected to the highly proliferative sympathetic phenotype suggested that pushing sympathoblasts into a chromaffin-like state may offer an interesting therapeutic strategy for neuroblastoma” which we will discuss below.

To address the anomalous position of cells in Cluster 6, we employed MultistageOT to force a differentiation trajectory originating from this cluster toward chromatin or sympathoblast fates, mirroring the normal dataset branching trajectory (Supplementary figure 5B). This approach generated new cell types (Supplementary figure 5C) and a time-dependent matrix for CardamomOT. However, the simulation results were quite revealing: the resulting UMAP displayed no discernible structure (Supplementary figure 5D). Investigation revealed that the simulated protein dynamics were completely flat (Supplementary figure 5G), lacking the temporal variation observed in the previous analysis (Figure 6F and G). This indicates that the forced trajectory was perpendicular to the true biological differentiation process. Crucially, this failure demonstrates that CardamomOT, while mathematically capable of processing any input, cannot generate meaningful biological dynamics if the underlying process is not structurally aligned with the data. The algorithm faithfully reflects the absence of a relevant dynamical trajectory in the input.

Assuming a differentiation sequence starting with chromaffin cells and ending with sympathoblasts, we proposed a framework through which cells could be organized, genes could be selected, and a 85 genes-based GRN could be inferred.

We would like to emphasize that contrary to most, if not all, GRN inference algorithms our gene selection procedure is completely blind to the function of the selected genes and relies purely on the time-dependent evolution of their mRNA distributions (see “Limitation of the study”). This implies that many genes selected are not transcription factors (TFs), a very clear difference from most GRN inference, where regulatory proteins within a GRN are restricted to TFs, as in ([70]; [71]; [72]). It is nevertheless of interest that the most densely connected node was *TCF4*, a basic Helix-Loop-Helix transcription factor that was recently shown to be directly involved in promoting neuroblastoma growth ([73], [74]).

The analysis of the subset of genes connected by specific dynamical motifs showed the existence of a group of genes, densely connected through various motifs like self-reinforcing loops, toggle switches FFLs and IFFLs. IFFLs have been demonstrated to be important for developmental processes like the generation of striped gene expression pattern ([55]) through morphogen gradients interpretation ([75]), or the fine tuning in cell-cycle progression ([76]). A careful examination of the impact of the different motifs on the fine-tuned gene expression pattern would require some systematic efforts to alter interactions values and documents the impact on the overall gene expression pattern, a task beyond the scope of the present work.

In order to validate our *in silico* approach, we tested the effect of *TCF4* inhibition. This could be in principle be carried out through the knock-out of *TCF4* as exemplified in ([58]) and by verifying that we indeed observe the predicted reduction in the amount of proliferating late sympathoblasts. Such a drive of sympathoblasts toward a chromaffin-like state has been proposed to represent a promising therapy strategy ([12]). This could be verified in our case by reinjecting mice with either WT or *TCF4* KO tumoroids and verify if those KO cells indeed have a reduced ability to generate tumors *in vivo*. A much faster and cheaper alternative was to use specific drugs, since *TCF4* gene-regulating activity can be targeted indirectly by using bromodomain and extra-terminal domain inhibitors (BETis), like JQ1, known to disrupt a TCF4-dependent super-enhancers ([59]). Furthermore, JQ1has been shown to inhibit human neuroblastoma cell lines growth and to induce their differentiation ([60]).

We therefore treated our PDTs with JQ1 and verified that:

- *TCF4* activity inhibition had a significant impact on PDTs growth
- *TCF4* activity inhibition had a highly significant impact on transcriptomic status of our PDTs, with hundreds of genes affected, both up- and down-regulated.
- JQ1-affected genes were enriched in functions not pertaining to cell division.
- *TCF4* activity inhibition led to a decrease in the amount of late sympathoblasts, mostly in favor of early sympathoblasts

The *PLK1* gene is another promising candidate as its *in silico* perturbation significantly alters cell type composition (see Figure 7A). Its expression is inversely correlated with neuroblastoma patient survival ([77]; [78]). This kinase serves as a key mitotic target and a neuroblastoma biomarker that enhances MYC stability, creating a feed-forward loop often upregulated in *MYC*-amplified tumors ([79]; [80]; [81]; [77]). Consequently, *PLK1* inhibitors like HMN-214, which have been validated to reduce neuroblastoma 3D spheroid tumor mass and growth ([78]) could be tested on our PDTs.

Finally, a combination of HMN-214 together with JQ1 should allow lower, non-toxic doses to be used for each drug, and be more efficient as predicted *in silico* (Figure 7B), and preventing or delaying the appearance of known resistance to JQ1 ([62]).

The results we obtained through JQ1 treatment support the idea that “pushing sympathoblasts into a chromaffin-like state may offer an interesting therapeutic strategy for neuroblastoma” ([12]). Of course, this whole study relies upon one single patient. The question as to whether or not the results we obtained can be relevant for more patients is quite open, especially given the remarkable heterogeneity of neuroblastomas ([69]). We probed the Childhood Solid Tumor Network (CSTN) Data Portal which indicated that all of the tested neuroblastoma cells including primary cells from OPDX from 4 different patients, were indeed sensitive to JQ1 addition (Supplementary figure 7).

Heterogeneity is of essence when it comes to neuroblastoma. We found observable heterogeneity across the various cultures we monitored (data not shown). This heterogeneity stems primarily from well-documented inter-patient variability, but results as well from a variability associated with PDX development in different mice and the establishment phases of tumoroids. This biological reality must be integrated into how we conceptualize anti-cancer therapies to ensure effective treatment and prevent patient relapse. The bioinformatic and biomathematical approaches we present here represent a significant asset for addressing this heterogeneity and to highlight the representativeness of cell types present in neuroblastomas from different patients. Our study is thus pioneering in describing the dynamics between chromaffin cells and sympathoblasts in human, opening the perspective of personalized treatments for neuroblastoma. The proposed systems biology approach, coupling *in silico* predictions to experimental validation should be applicable to all biological processes involving differentiation dynamics.

We finally have shown that our PDTs do harbor a very marked spatial structure regarding the repartition between stem and progenitor cells ([28]). In order to assess whether or not the chromaffin to sympathoblasts trajectory we describe in the present paper leads to some kind of similar spatial structure in our PDTs or not, we are planning to generate and analyze data from single-cell transcriptomics studies using the BMKManu S3000 platform ([82]). Analyzing such spatial data would also be an opportunity to add cell-cell interactions in our modelling scheme. A recent proposal based upon a coupled system of partial and ordinary differential equations ([83]) could be an inspiration on how to couple our GRN with ligand concentrations.

### Limitations of the study

As we mentioned in ([21]), the ideal case study for CardamomOT is “any biological process in which time-stamped single-cell transcriptomic data are obtained after applying a given stimulus”. We could add that a predefined gene set and predefined cell types would also render the procedure smoother. Any departure from that scenario requires the application of *ad hoc* procedures, that are discussed below.

A specific “time” definition has to be proposed when the whole differentiation process is sampled at once. In ([52]) we defined overlapping windows, arranged along a “stemness axis”. We have shown in ([21]) that the sampling has to be sufficient in regard to the process. This led us to propose 5 time-points and an overall duration of the process of 50 hours. One nice validation of this choice of time is that it correctly captures the evolution of mRNA distribution (see Figure 6C), and that it allows for sufficient amount of time to also see variation in the amount of predicted protein amounts (see Figure 6F and G). Such a protein variation would not be seen for an inadequate temporal resolution (say 5 minutes windows for 25 minutes time). Interestingly, the output of the model seems quite robust to the choice of number of time points (from 3 to 10; see Supplementary figure 4).

A gene selection procedure has to be proposed. This can be a small list of genes of interest like in ([84]), a literature-based list like in ([21]), or a double selection procedure (using both Kantorovich distances and entropy) as proposed here. Our gene selection method relies on an arbitrary threshold for the number of genes, which may exclude biologically relevant ones. Unlike most GRN inference approaches, our selection is blind to gene function and based solely on the time-dependent evolution of mRNA distributions. This means many selected genes are not transcription factors (TFs), a key difference from traditional methods that focus on TFs (e.g. [70]; [71]; [72]). Additionally, our approach captures only genes with observable changes in mRNA distribution within the experimental time frame, likely missing faster interactions (e.g., protein phosphorylation). Thus, the edges in our GRN should be interpreted with caution, since they may represent indirect interactions. Their actual molecular mechanisms would require further validation with complementary techniques. Furthermore, the non pre-selection of gene function leads to the selection of long non coding RNAs (lncRNAs), where the “protein produced by a lncRNA” has no biological meaning *per se* but represents the influence that this lncRNA might have on target genes. It might be seen as a “metavariable” summing up undefined molecular processes. We nevertheless think it is quite relevant to keep those lncRNAs in our network, especially in the light of their known importance in neuroblastoma ([47]). We also noted that two of our lncRNAs (*MEG3* ([49]) and *MIAT* ([50])) have been shown to be functionally involved in neuroblastomas. The realism of our inferred GRN should therefore be nuanced: we do use a mechanistic model to describe gene expression, but this model has to be abstracted to account for interaction that may not be direct and may also needs to be abstracted to describes the influence of genes not abiding by classical assumptions.

The definition of cell types is not a requirement *per se* in our GRN inference procedure. This is nevertheless needed when one tries to define the effect of a perturbation at the cellular level. Here we tried to define NB cell types based upon known signatures from normal cell types. The variations of those signatures are quite continuous along the proposed differentiation sequence (see Figure 1E and 1F). This led us to propose a definition of cell types based upon clustering and organized along the proposed differentiation sequence. Initially, cell types were defined by aggregating Leiden-computed clusters. Using Louvain-computed clusters instead seems to make only small differences (see Supplementary figures 3 and 4).

We would like to stress out that cell types were defined by aggregating Leiden-computed clusters, whereas the time-points have been defined by slicing through the velocity pseudotime. The ability to accurately reproduce cell type proportions suggests that these two choices are well-suited.

Finally, we would like to propose that all those choices define a “window” through which one observe a complex mechanism. Varying the size and location of these window (through different parameter choice combinations), different part of the outside world will be viewed. Those views are true but fundamentally incomplete. The experimental validation is in our mind the best to make sure the view from a given window is real.

---

**Supplementary figure 1:**
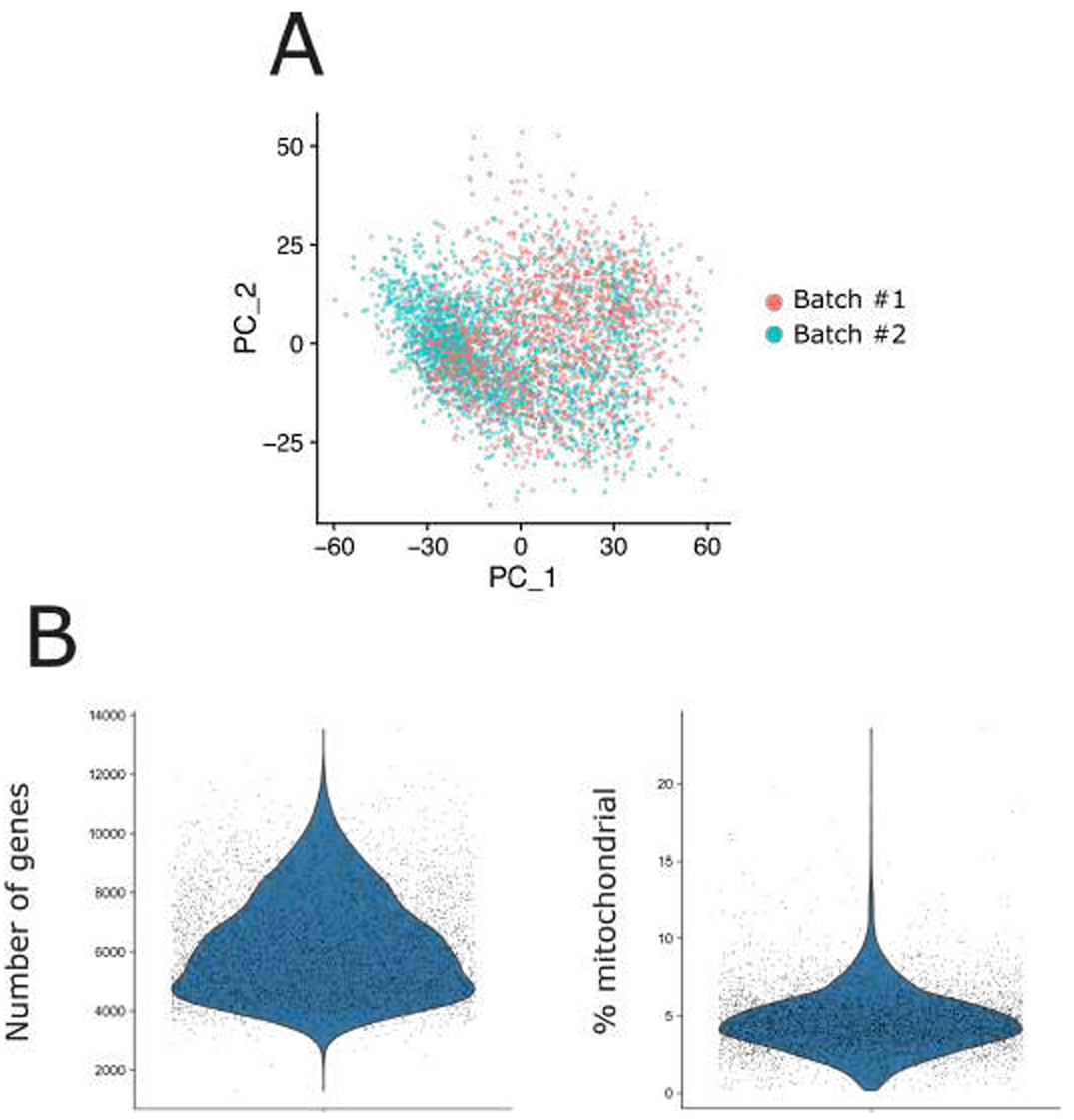
Low-level analysis of the dataset. A: PCA analysis of the two batches. B: QC metrics displayed as a violin plot, displaying the number of genes and the percent of mitochondrial genes per cell.

**Supplementary figure 2:**
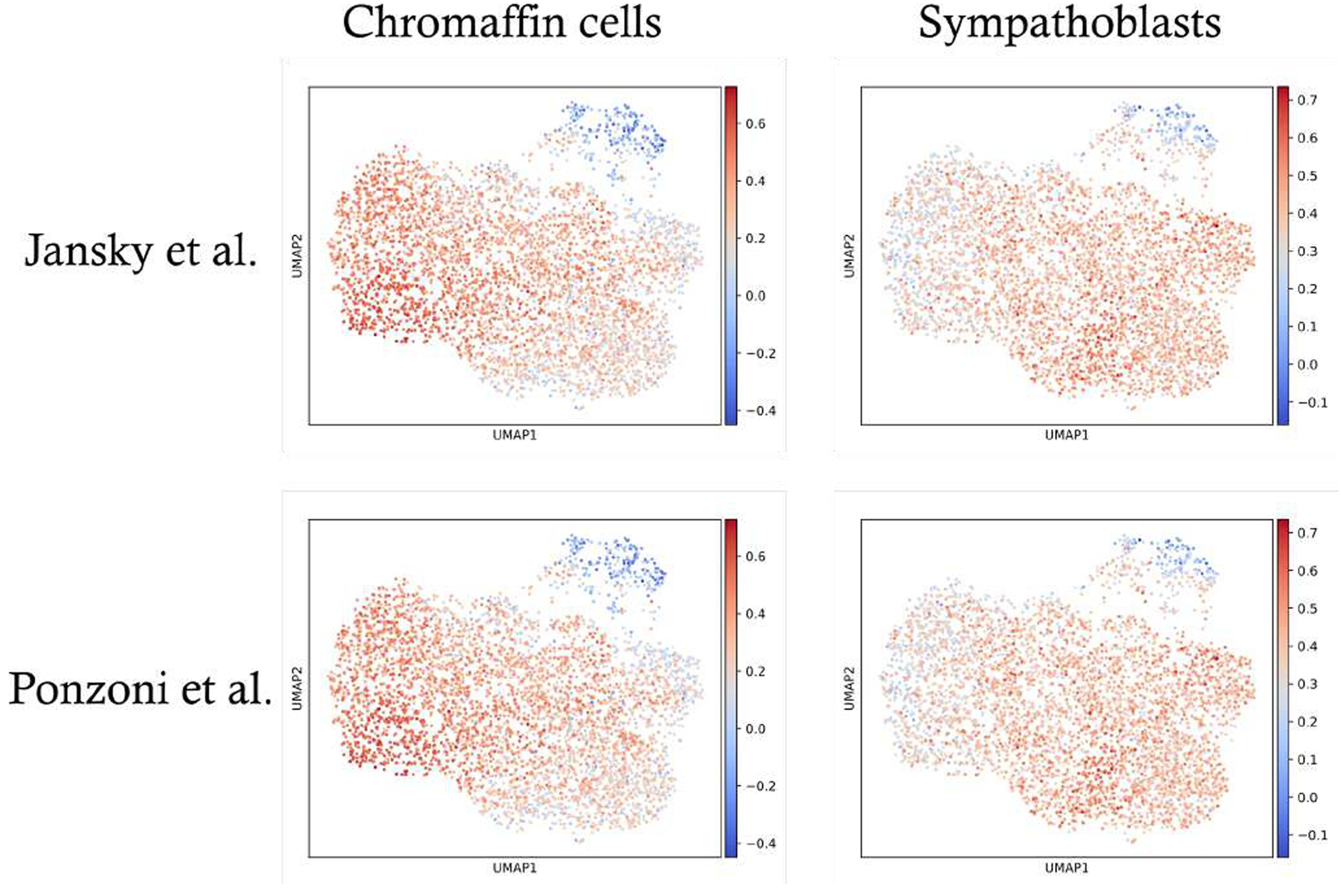
Expression level of two published gene signatures ([11] and [7]; see material and method for the gene lists) to identify the presence of chromaffin cells and sympathoblasts.

**Supplementary figure 3:**
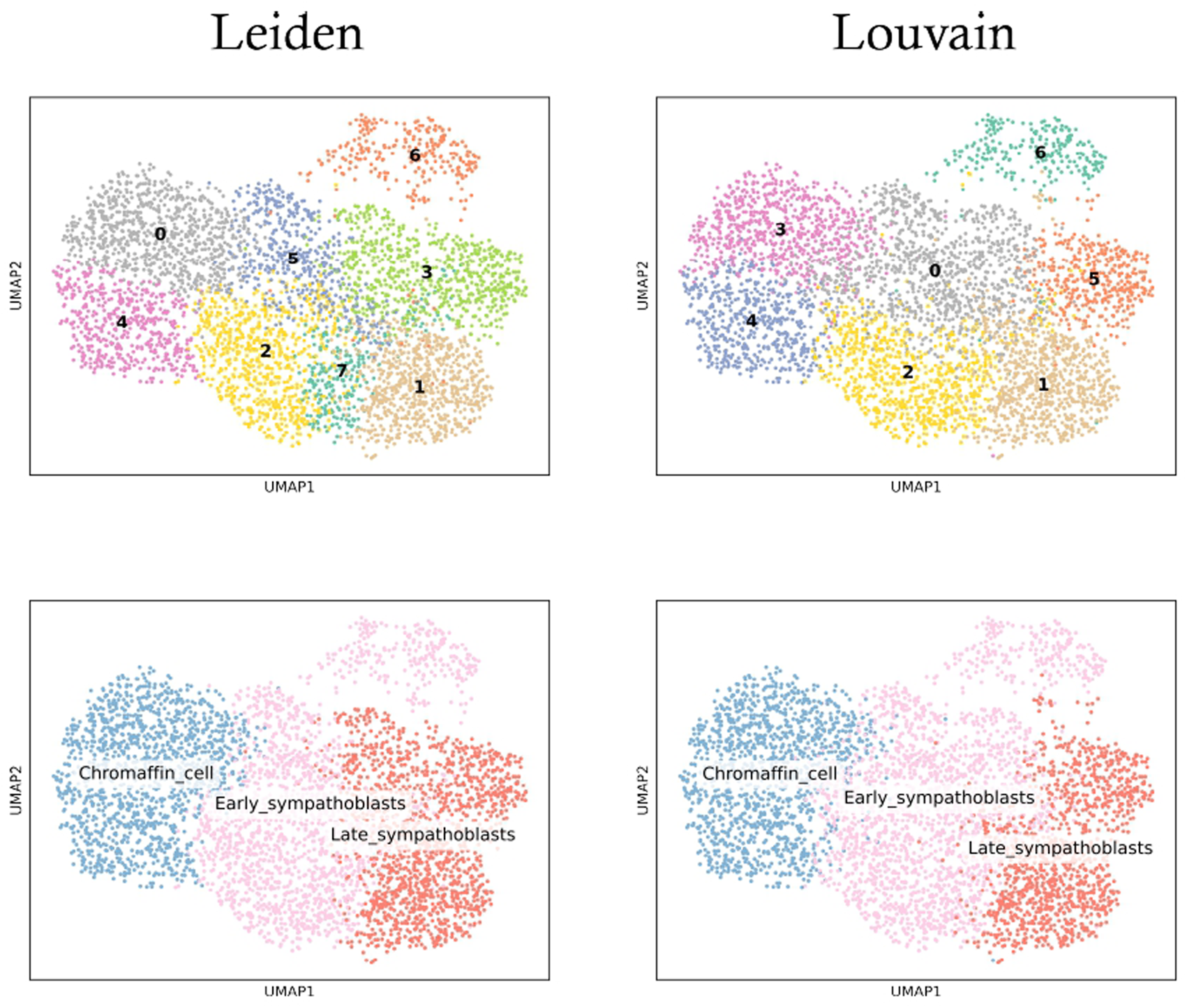
The result of the Leiden clustering is shown on the left, and that of the Louvain clustering on the right.The bottom shows the cell type definition made by aggregating the above clusters. The adjusted rand index is of 0.60 for the clusters and of 0.66 for the cell types definition.

**Supplementary figure 4:**
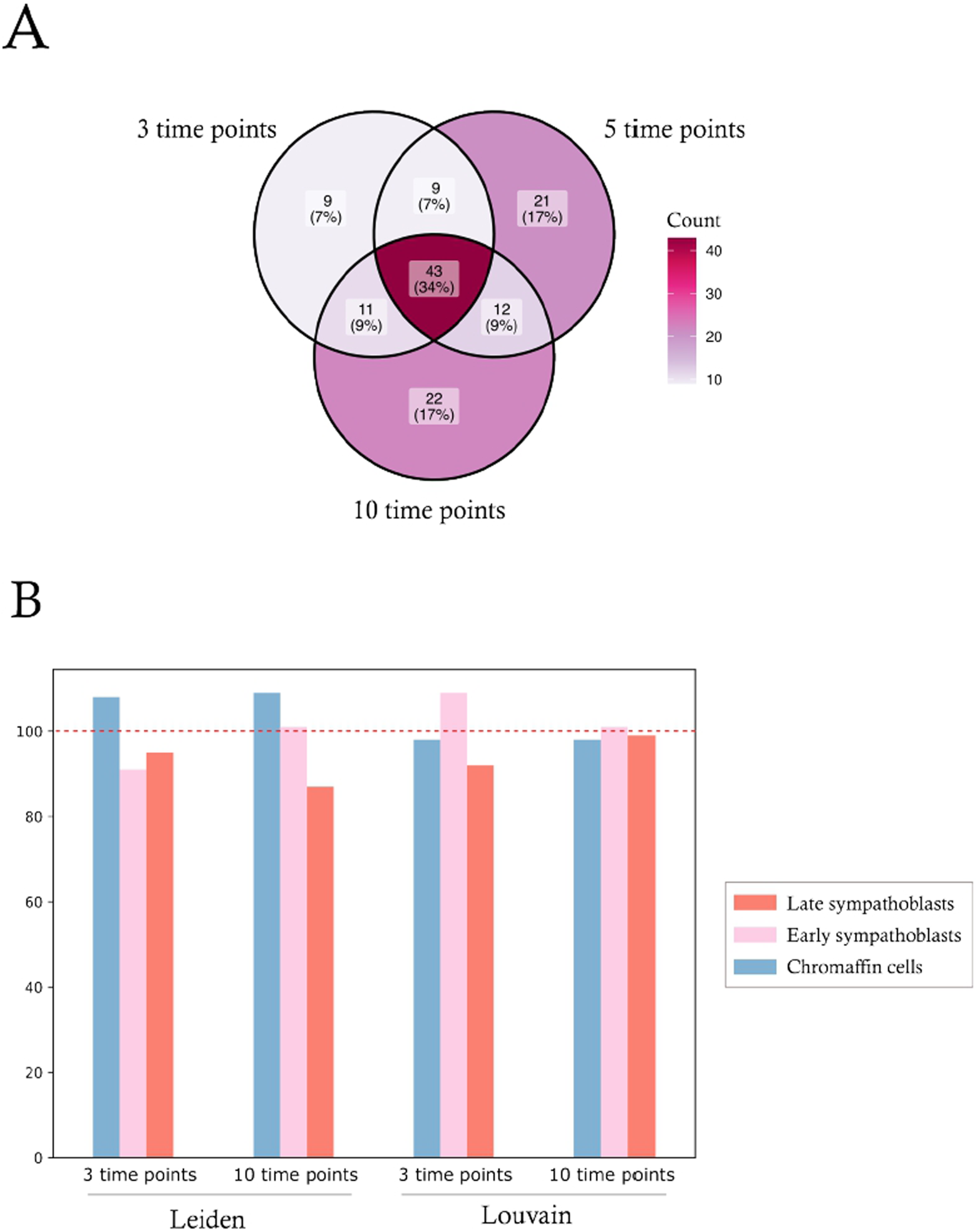
A: overlap between the genes lists made when considering 3, 5 or 10 time points along the velocity pseudotime. B: ability of the GRN inferred in those 3 cases to generate the expected cell types (experimental), either defined through Leiden or Louvain clustering (see Supplementary figure 3 above). Shown is the comparison between the number of cells computed for each cell type compared to the repartition observed in the 5-time points case: when using a GRN inferred using 3 time points, this GRN tends to slightly overestimate the number of chromaffin cells defined using Leiden clustering and to slightly underestimate the number of chromaffin cells defined using Louvain clustering.

**Supplementary figure 5:**
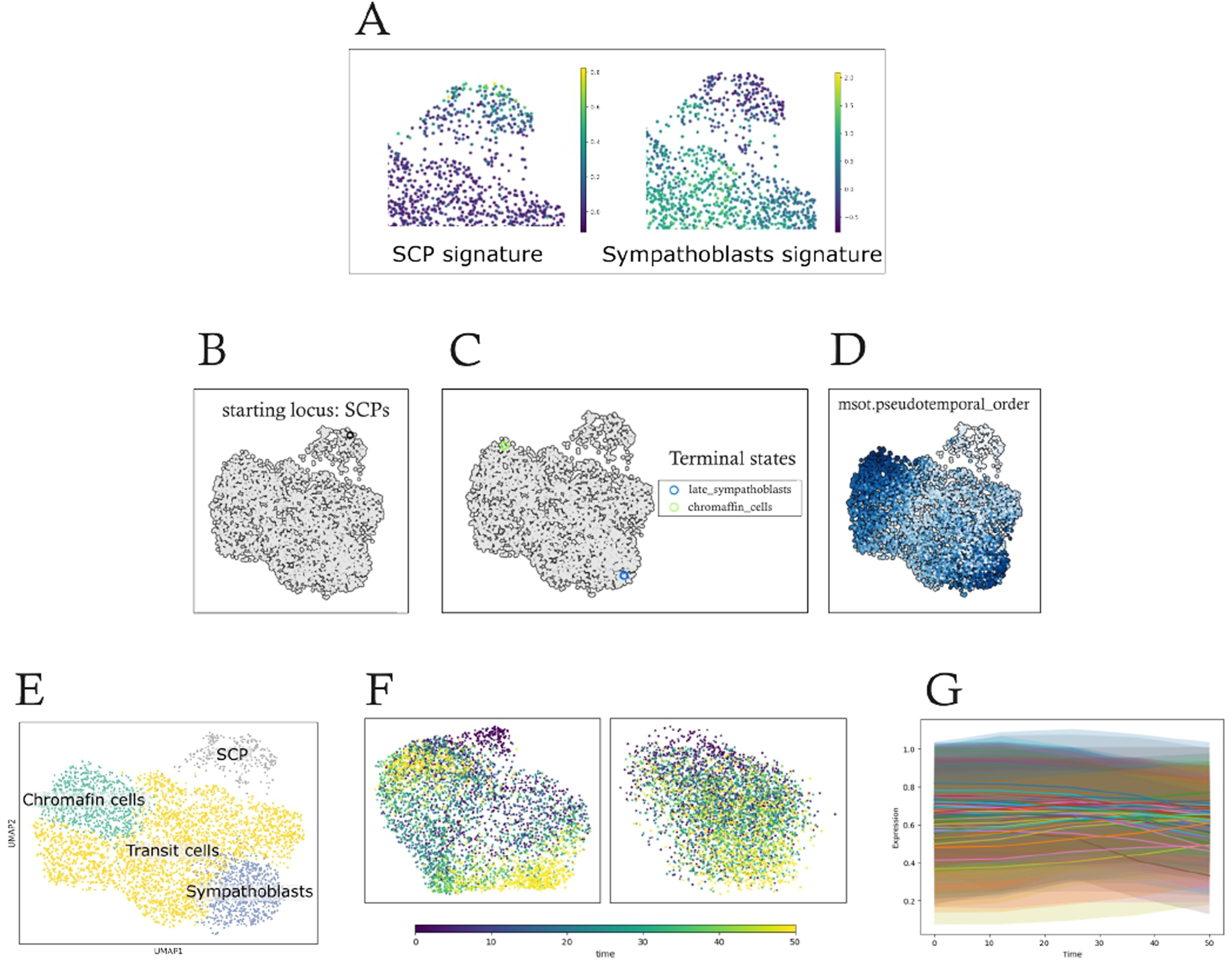
Cluster 6 exploration. A: zoom on cells from the cluster 6 color coded according to an SCP signature or a sympathoblasts signature. B to D: MultistageOT analysis. We first defined a starting locus (B), then terminal states (C) and ran MultistageOT to compute a pseudotemporal_order (D). E: cell type definition destined to reproduce the bifurcation fork seen in the normal differentiation sequence when imposing cluster 6 as the start; F: UMAP representation of the dataset used for GRN inference (left) and of the dataset obtained through GRN simulation (right); G: time-dependent evolution of the mean+/-SD of all proteins expression.

**Supplementary figure 6:**
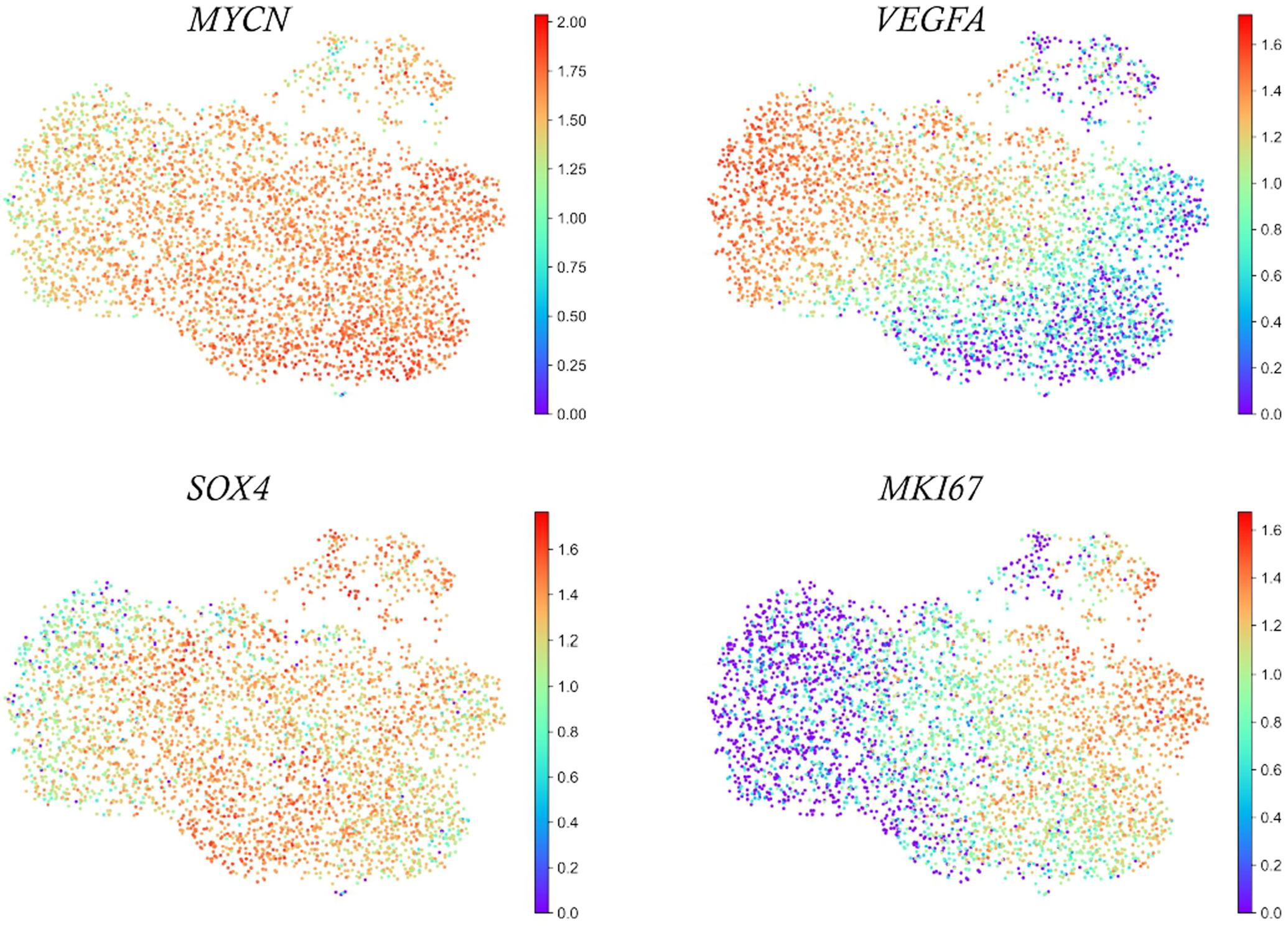
UMAP representation where each cell is color-coded according to its gene expression level.

**Supplementary figure 7:**
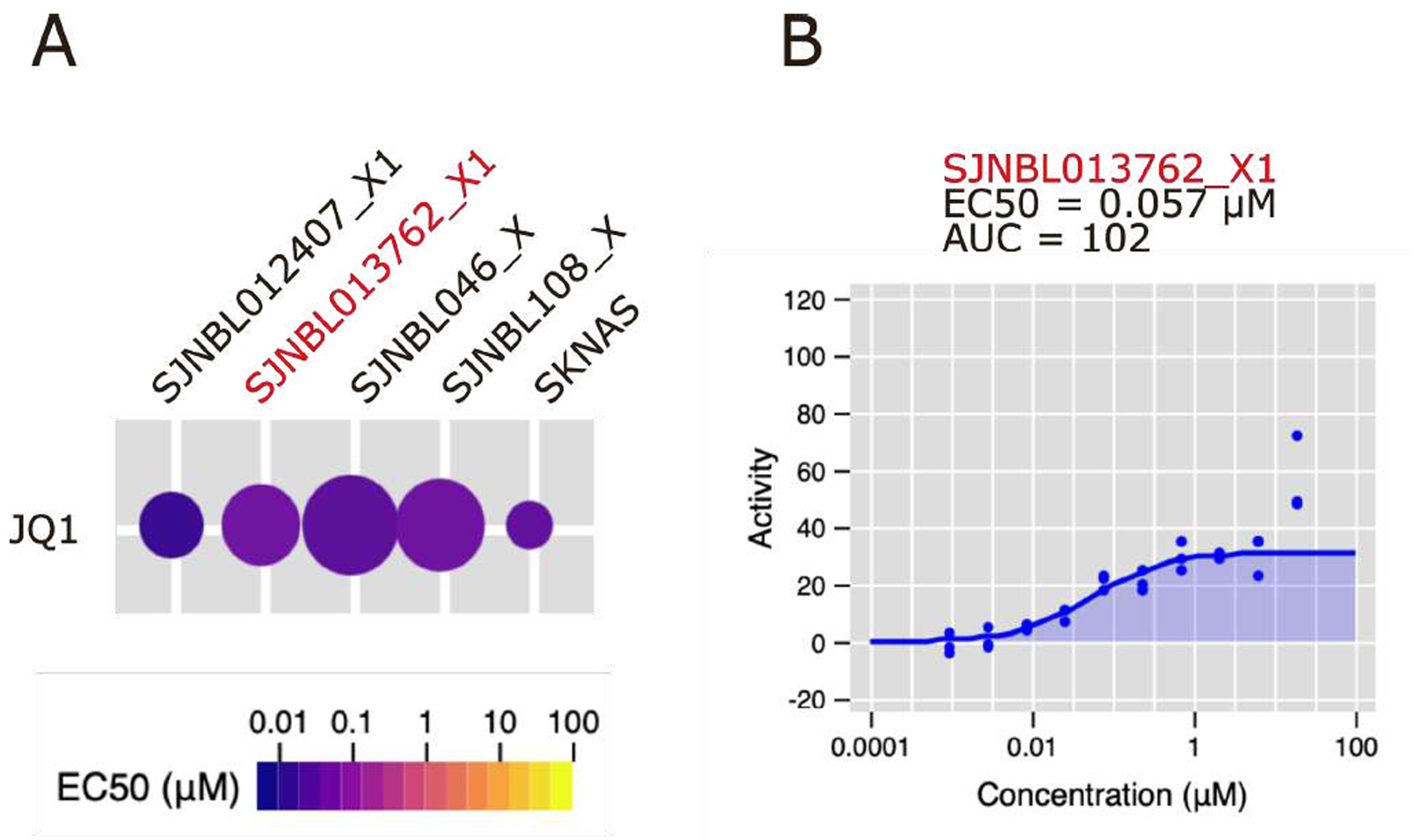
Neuroblastoma cells sensitivity to JQ1 as provided by the Childhood Solid Tumor Network (CSTN; https://braid.stjude.org/masttour/) Data Portal. A: The primary heatmap indicates the potency of the compound (EC50 in μM) with color, the circle size indicating the efficacy (area under the curve, or AUC). The OPDX we used in this study is highlighted in red. _X indicates primary cells derived from OPDX. SKNAS is an established NB cell line. B: Dose response for the SJNBL013762_X1 primary cells.

**Supplementary Table 1:** the list of the 85 genes kept for constructing our GRN.

|  |  |  |  |  |  |  |
| --- | --- | --- | --- | --- | --- | --- |
| *ACAT2* | *CHAF1A* | *GATA2* | *KPNA2* | *MT.CO1* | *PLK1* | *TOP2A* |
| *ANP32E* | *CHGA* | *GNAS* | *MAB21L1* | *MT.CO2* | *PRR11* | *TPX2* |
| *ARL6IP1* | *CRABP1* | *H1.0* | *MCM2* | *MT.CO3* | *PTTG1* | *TUBA1B* |
| *ASPM* | *CYGB* | *H19* | *MCM3* | *MT.CYB* | *RAD21* | *UBE2S* |
| *BIRC5* | *DDIT4* | *H4C3* | *MCM4* | *MT.ND3* | *RGS4* | *VEGFA* |
| *BNIP3* | *DHFR* | *HMGB1* | *MCM6* | *MTATP6P1* | *RRM2* | *VIM* |
| *BRCA1* | *DLK1* | *HMGB2* | *MCM7* | *MTND1P23* | *SMC4* | *ZFAS1* |
| *CALM2* | *DNAJB1* | *HMGCS1* | *MDK* | *MYCN* | *SNHG7* |  |
| *CCNB1* | *DUT* | *HSPA1B* | *MEG3* | *NEFM* | *STMN4* |  |
| *CDC25A* | *E2F1* | *IGFBP2* | *MIAT* | *NRP2* | *TAF1D* |  |
| *CDKN1C* | *FAM49A* | *IGFBP5* | *MKI67* | *PCNA* | *TANC2* |  |
| *CENPE* | *FEN1* | *KDM5B* | *MT.ATP6* | *PFKP* | *TCF4* |  |
| *CENPF* | *GAS5* | *KIF21A* | *MT.ATP8* | *PHOX2B* | *TH* |  |

## Ethics approval and consent to participate

Frozen cells from orthotopic patient-derived xenograft were obtained from the biorepository of St Jude Children’s Research Hospital (St. Jude Children’s Research Hospital, 262 Danny Thomas Pl, Memphis, TN 38105, USA) at https://www.stjude.org/research/why-st-jude/shared-resources/biorepository.html. Formal patient consent was obtained by St Jude Children’s Research Hospital as indicated in ([85]) under the Molecular Analysis of Solid Tumors (MAST) protocol (ClinicalTrials.gov ID NCT01050296, https://clinicaltrials.gov/ct2/show/NCT01050296; Clinical trial number: not applicable.).

## Consent for publication

All authors approved the publication.

## Authors’ contributions

Conceptualization, O.G. and S.G.-G.;

Data Generation, C.K. and E.V.;

Methodology, O.G and E.V.;

Project administration, O.G.;

Software, O.G. and F.P.;

Supervision, O.G.;

Writing – original draft, O.G.;

Writing – review & editing, All.

## Acknowledgements

We are deeply indebted toward Elias Ventre (INRIA) for his invaluable help in using CardamomOT and during the revision phase of the manuscript. We thank Tess Chilliet (YCBL student) for an early and relevant exploration of our dataset and Christophe Arpin (CRCL) for sharing unpublished material regarding gene selection procedure.

We thank the computational center of IN2P3 (Villeurbanne/France), where the computations were performed. We also thank the BioSyL Federation and LabEx Ecofect (ANR-11-LABX-0048) of the University of Lyon for inspiring scientific events.

Part of the code was debugged using Le Chat (https://chat.mistral.ai/) or Euria (https://www.infomaniak.com/fr/euria). IAG was also sometimes used for rephrasing in proper English.

This work was not financed by the Association Pour la Recherche Contre le Cancer (ARC).

## Software availability

CardamomOT is available from https://cardamomot.readthedocs.io/en/latest/index.html.

A git depot is available at: https://github.com/eliasventre/CardamomOT. We used the *plos-cb-2026* version.

Three scripts that can be used to prepare and analyze a CardamomOT-ready matrix from fastq files are available from: https://github.com/ogandril/From_fastq_to_CardamomOT

The scripts used for launching CardamomOT on a distant server are available at https://github.com/ogandril/Launching_Cardamom. ktest is available from https://github.com/LMJL-Alea/ktest. License: BSD 3-Clause License

## Data availability

Both raw sequences and a processed version can be found at:

- https://www.ebi.ac.uk/biostudies/arrayexpress/studies/E-MTAB-16741 for the original PDTs RNAseq experiments.
- https://www.ebi.ac.uk/biostudies/arrayexpress/studies/E-MTAB-16963 for the JQ1 treatment experiment.

The dataset can be explored via: https://bioshiny.ens-lyon.fr/public/app/ogsinglecell

## Material and methods

### Tumoroids generation and culture

Tumoroids were grown as previously described ([28]). Briefly, frozen cells from orthotopic patient-derived xenograft (OPDX; [85]) sample number: SJNBL013762_X1 (https://cstn.stjude.cloud/sample/SJNBL013762_X1#) were provided by the St. Jude Children’s Research Hospital through the Childhood Solid Tumor Network. This OPDX was derived from a tumor removed from a 15 month-old boy, with adrenal primary tumor location and bone marrow metastasis, which harbored known molecular alterations consisting in *MYCN* gene amplification and *alk* mutation. The study had all necessary regulatory approvals and informed consents are available for all patients (see “ Ethics approval and consent to participate”).

Patient-derived xenograft (PDX) were generated according to [85]: 10^6^ thawed OPDX cells were injected subcutaneously after embedding in 100 µL matrigel (Corning™ Matrice Matrigel™ ref 356234) in 6 weeks old NSG-NOD.Cg-Prkdcscid Il2rgtm1Wjl/SzJ (Charles River ref. 614) female mice. The mice were housed in sterilized filter-topped cages and maintained in the P-PAC pathogen-free animal facility (D 69 388 0202; Cancer Research Center of Lyon; CRCL). All animal studies were performed in strict compliance with relevant guidelines validated by the local Animal Ethic Evaluation Committee (C2EA-15) and authorized by the French Ministry of Education and Research (Authorization APAFIS\#28836).

Fresh tissues from tumor developed in NSG-NOD mice were mechanically minced into small pieces and then digested by using the gentleMACS™ Dissociator (Miltenyi Biotec) with Tumor Dissociation Kit, human (ref 130-095-929) in C-tube program SOFT 37_h_TDK_1. After 61 min incubation at 37 °C cells were collected and purified with Debris Removal Solution (ref 130-109-398), Red Blood Cell Lysis Solution (ref 130-094-183) and Mouse Cell Depletion Kit (ref 130-104-694). 4.10^4^ cells were seeded after embedding in 10 µL matrigel and then placed in 96-well ULA plates (BIOFLOAT™ 96-well U bottom - FaCellitate) in 100 µL culture medium based on previously described neuroblastoma tumor-initiating cell lines and PDT derivation protocols ([86]; [27]) which consists of advanced DMEM/F-12 medium (Gibco), 1X B-27 supplement without vitamin A (Gibco), 1X N2 supplement (Gibco), 1x Penicillin-Streptomycin (Gibco), 40 ng/mL hFGF-b (Peprotech), 20 ng/mL hEGF (Peprotech), 10 ng/mL hPDGF-AA (Peprotech), 200 ng/mL hIGF1 (Peprotech) and 10 ng/mL hPDGF-BB (Peprotech).

Cultures were then trypsinized using TrypLE Express Enzyme (ThermoFisher Scientific) and split without embedding in the same medium and the same 96-well ULA plates. Cultures were supplemented after 7 days by the addition of 50 µL of medium, and then the medium was renewed by half twice a week. PDTs were split by trypsinization every 15-19 days when reaching a diameter of about 1 mm.

The neuroblastoma nature of the growing tumoroids was confirmed by immunohistochemistry performed with the following antibodies: anti-CD56, anti-PHOX2B and anti-SYP (SYNAPTOPHYSIN) (see ([28]) for the last one).

All experiments were performed from a unique stock of tumoroid frozen at passage 8. For freezing, 20 tumoroids (about 5.10^4^ cells each) were harvested and centrifuged 5 min at 1500 rpm. The pellet was gently resuspended by pipetting up and down in 200 µL STEMCELL Technologies CryoStor™ CS10 at room temperature and transferred into cryovials. Those were placed into Mr. Frosty Freezing Container and transferred to −80 °C for at least 2 days before being stored at −150 °C for long term storage.

For thawing, cryovials were rapidly warmed in a 37°C water-bath until only a small ice crystal remained, and then placed on ice. The cell suspension was transferred from the cryovial into a tube with 10 mL RPMI medium at room temperature, and centrifuged for 10 min at 750 rpm. The pellet was resuspended in pre-warmed tumoroid culture medium supplemented with 10 µM Y-27632 (ROCK inhibitor, STEMCELL Technologies). Thawed tumoroids were seeded in 10 wells of a 96-wells plate and incubated at 37 °C, 5 % CO₂. The next day the medium containing the ROCK inhibitor was replaced by the standard tumoroid culture medium. 4 days later the tumoroids were splitted at 15000 cell per well in a 96 well-plate and supplemented after 7 days by the addition of 50 µL of medium. Cultures were then maintained using the same routine: supplemented after 7 days by the addition of 50 µL of medium, splitting on day 14 and reseeding them at 15,000 cells per well.

### Data generation and filtering

scRNAseq was performed using RevGel-seq ([29]) with few modification from the Scipio bioscience protocol.

Briefly, PDTs were trypsinized to obtain a single-cell suspension, which were then chemically labeled to enable interaction with barcoded beads. For this, cells were incubated with a bifunctional chemical linker (polyA at one extremity and a hydrophobic moiety at the other extremity) in DPBS for 5 min. Labeled cells were resuspended at a concentration of 300 cells/µL, yielding a total input of 15,000 cells.

To form cell-bead complexes, 50 µL of the barcoded bead suspension was pipetted into the bottom of a gelation tube, followed by the addition of 50 µL of the labeled cell suspension. The mixture was immediately homogenized by pipetting up and down five times at a rate of 90 rpm to facilitate coupling through collisions. The excess of barcoded beads ensured that each bead bound to no more than one cell. The cell-bead suspension was then diluted with 2.4 mL of hydrogel solution and homogenized by gently rotating the gelation tube. The gelation piston was slowly inserted into the tube while keeping it vertical. The device was incubated on ice for 20 minutes to complete gelation and immobilize the cell-bead complexes.

After removing the piston, cells were lysed with 7.5 mL of lysis buffer. At this stage, samples could be stored at -80°C. Incubation at room temperature with orbital shaking allowed cell lysis and local release of RNA molecules, which hybridized to the poly-T extremities of the capture beads. Gel dissolution was achieved by adding 3.75 mL of degelation buffer and vortexing for 5 minutes, resulting in a mixture of RNA hybridized to the barcoded beads.

Reverse Transcription (RT) and PCR were performed on the bead-bound polyA tailed RNA. Reverse transcription was carried out using the *Maxima H Minus RT kit* (ThermoFisher, EP0751) for 10 min at 25°C followed by 90 min at 42°C. A 50-minute incubation at 37°C with Endonuclease I eliminated single-stranded sequences and remaining oligos. Alkaline denaturation with 50 µL of 0.1 M NaOH for 5 minutes removed RNA, leaving cDNA bound to the beads. The second strand was synthesized using *S3 Supermix* (1.1× RT buffer, 1.3 mM each dNTP, 12% PEG8000, 13 µM second strand primer) during an 1-hour incubation at 37°C. PCR amplification was performed with *KAPA HiFi HotStart Readymix (*Roche, 07958935001). The cDNA was purified by pooling the PCR products and using 0.6x volume of *SPRIselect reagent* (Beckman Coulter, B23317).

The quality of the purified cDNA amplicons was assessed using the *Agilent TapeStation HSD5000*, and quantification was performed with the *Qubit dsDNA HS Assay Kit (*Invitrogen). 20-30ng of purified cDNA were processed using the *Illumina Nextera XT Library Preparation Kit* (Illumina, FC-131-1024), with a 5-minute fragmentation step at 37°C. A custom library prep primer (provided by Scipio) was used for the P5/read1 side of the library amplification PCR. 10 cycles of PCR were performed to generate a library ready for sequencing with the *Illumina NextSeq500* platform (*NextSeq High Output Flow Cell*, paired-end, 25/0/0/67).

The raw sequence data was mapped to the human genome (GRCh38) using STARsolo ([87]) run in the Velocyto mode after creating the whithelist containing the cell tags from the sequences. Genes were identified using the GRCh38.99.chr.gtf file.

The resulting .mtx matrices were opened using Seurat 5.4.0 ([88]) and combined for comparison purposes. Since both datasets were clearly not separated by PCA (Supplementary figure 1), we decided to aggregate them into a unique dataset.

Aggregation was performed using scanpy ([36]). The merged dataset was filtered to keep only genes having a non null expression value in 6 cells, and only cells with a least 100 genes expressed. This resulted in a 30034 genes x 4345 cells expression matrix that was normalized using median count depth normalization with log1p transformation.

UMAP was computed on 2000 HVG selected using Seurat FindVariableGenes function. Very similar results were obtained after gene selection using GeneCover ([89]) or mixHVG ([90]; not shown) The distribution in the number of genes and the percent of mitochondrial DNA is shown in Supplementary figure 1B.

Gene Ontology enrichment was performed using ClusterProfiler ([51]).

### Tumoroids treatment

PDTs were treated with 10^-6^ M JQ1 (ref 126331-10212-1, Tebu Bio) for 24h as described ([91]) and analyzed by RevGel-seq (see upper). We obtained 2201 cells and 30034 genes for the control condition and 1532 cells and 30034 genes for the JQ1-treated condition. Data analysis was performed as described upper. On the other hand, treatment was done for 48 hours for assessing JQ1 impact on cell number, and cell numbers were counted at 24 and 48 hours.

### Signature genes

We used three gene signatures for Chromaffin cells and sympathoblasts:

1. We used by default the one defined in ([9]) as: Chromaffin cells: *TH, PENK, CHGA, CHGB, DGKK, DLK1* and *EPAS1*. Sympathoblasts: *ISL1, NEFM, NEFL, STMN2, ELAVL2, ELAVL4, CARTPT, PRPH, LIX1, CPLX1* and *EGLN3*.
2. We also used the signature defined in ([7]) as: Chromaffin cells: *CHGA, CHGB, TH, PENK, DBH, SOX10, PLP1, RET, INSM1, PNMT, ASCL1, PHOX2B and CDKN1C Sympathoblasts: RET, ASCL1, PHOX2B, HAND2, GATA3, TH, ISL1, ALK, MEIS2, NPY, PRPH, NEFM, STMN2, SYN3, CCND1, MKI67* and *TOP2A*
3. The third signature used was derived from ([11]): Chromaffin cells: *DBH, TH, CHGA, DDC and PNMT* Sympathoblasts: *MKI67, TOP2A, NEFM, GAP43, STMN2, ISL1, ALK, SYN3* and *IL7* For Schwann cell precursors (SCPs) we used the same signature than ([52]): *PLP1, FOXD3, FABP7, S100B, NGFR, ERBB3, MBP, MPZ, COL2A1, POSTN, MOXD1* and *GAS7*.

Signature intensity was computed with the score_genes function of scanpy, which computes the average of the normalized expressions of the signature genes.

### Cell cycle scores

We used the scvelo.tl.score_genes_cell_cycle function to calculates scores and assigns a cell cycle phase (G1, S, G2M) using the list of cell cycle genes defined in ([92]).

### Differential expression analysis

We used the sc.tl.rank_genes_groups function which computes a t-test and corrects for multiple testing using the Benjamini-Hochberg correction.

### Defining the GRN genes

To estimate the Shannon entropy, the best upper bound estimator (BUB) was computed as developed in ([93]). The R script for this estimation (BUB_discrete_entropy.R) is available on this OSF project: https://osf.io/9mcwg/. The python implementation is available in the https://github.com/ogandril/From_fastq_to_CardamomOT/Define_genes.ipynb file.

The Kantorovich distance was computed using the wasserstein_distance function from the scipy.stats package.

### CARDAMOM-based inference: general overview

CARDAMOM-based inference frames the GRN inference problem as calibrating a mechanistic model of gene expression using temporal snapshots of scRNA-seq data ([21]). In this model of gene expression called HARISSA ([94]), transcriptional bursting is captured by a two-state gene expression model where each gene stochastically switches between an inactive (OFF) state with no transcription and an active (ON) state with high transcription, so that mRNAs are produced in bursts whenever the gene is ON. The bursty version thus explains single-cell expression as the result of these random state switches, naturally generating non-Gaussian, overdispersed distributions (e.g. gamma/negative binomial) which reproduce the main statistical characteristics of the experimental data ([21]). Coupling the genes through interactions, the value of which has to be fitted, generates the GRN.

The dynamics start with a small number of genes being turned on by the stimulus. In our previous studies, the stimulus was explicitly identified as an exogenous medium change/addition ([95]; [21]). Here, it should be understood as a complex influence exerted by the cell environment, whether hormonal or through cell-cell interactions, which induces cell differentiation. It is needed to “set the GRN in motion” and push the cells out of their quasi-steady state. Its value is 0 at time 0 and 1 for all the consecutive time points.

### GRN inference: parameters

All computations were performed at the IN2P3 computing center (https://cc.in2p3.fr). The following parameters were used:

- SFT= 10 #Correspondance between velocity pseudotime and model time
- stim = 0.22 #Controls for the overall influence of the stimulus.
- m=0.75 #self.mean_forcing_em; defines the intensity of constraining the GRN via the mean of the marginals.

In order to mitigate run-to-run variability due to the stochasticity of the inference algorithm, we ran 10 inferences in parallel and computed a mean version of the GRN. For this, the mean value along the 10 repeats was computed for each interaction and kept. In order to assess the run-to-run variability, see Figure 10 in ([52]).

This mean version is the one displayed and analyzed in figures 4 and following. The simulations and perturbations experiments were carried out by forcing this version to be the GRN used by setting the following values in the base.py file:

- `self.hard_forcing_ref = True`
- `self.ref_constraint_pct = 0`
- `self.seuil_min_network = 0`
- `self.lambda_mlp = .5`
- `self.filter_network = 1`
- `self.seuil_min_network_intensity = 0.048`
- `self.seuil_min_network_variations = 0 and an inter_ref.csv file containing the reference GRN in the data folder.`

### Motifs exploration

Custom R scripts were developed to systematically traverse the network, identifying all feed-forward loops (FFLs) and incoherent feed-forward loops (IFFLs), and classifying them based on the algebraic signs of their three constituent interactions. A Monte Carlo randomization procedure (1,000 iterations) was then performed by shuffling interaction signs while preserving the network topology, enabling the calculation of Z-scores to determine if the observed motif frequencies significantly deviate from a random distribution.

### GRN simulation

The simulation was performed using the CardamomOT build-in version of HARISSA, a software for performing GRN simulations based on an underlying stochastic dynamical model driven by the transcriptional bursting phenomenon ([94]).

The dynamics of each gene is given by the following ordinary differential equations:

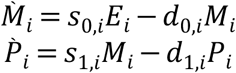

Where M_i_ is the number of mRNA molecules for gene i, P_i_ is the number of proteins for gene I, E_i_ is the state of the promoter, s_0,i_ is the mRNA synthesis rate, d_0,i_ is the mRNA degradation rate, s_1,i_ is the protein synthesis rate, and d_1,i_ is the protein degradation rate. E_i_ can randomly switch from 0 to 1 (activation) with a protein-dependent rate k_on,i_ (P) and from 1 to 0 (inactivation) with a constant rate k_off,i_.

We define the protein-dependent rate function k_on,i_ as in ([_94_]).

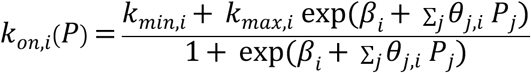

such that the promoter activation rate is between k_min,i_ and k_max,i_. The parameter *β_i_* represents the basal activity of gene i, and the parameter *θ*_j,*i*_ encodes the value of the j → i interaction (see below for its determination).

Because we simulated a mechanistic model, one needs to provide HARISSA with half-lives for both mRNAs (d_0,i_) and proteins (d_1,i_). Contrary to the early version of CardamomOT where those values had to be given from external sources, in the latest version we used, those values were directly inferred trough CardamomOT.

### *In silico* perturbations

KO were performed as described ([20])): the gene’s protein level is fixed at zero and the gene basal parameter βi is set to−1000. CardamomOT learns the identities of the three cell types using a random forest on the experimental dataset, and applies it to the model-generated dataset. The classifier was trained using the RandomForestClassifier function from Scikit-Learn with n_estimators=100 and labels were transferred using the predict function.

### ktest

ktest enables differential analysis within a non-linear kernel-based framework ([61]). This approach compares the distributions of gene expression or any molecular features through both global and feature-specific comparisons, and it can be extended to any measured single-cell feature. The code was run using default parameters. We computed the 4000 highly variable genes using scanpy `highly_variable_genes` function in each condition, and reduced the matrix to a set of shared 1533 HVGs.

### GO overrepresentation

GO overrepresentation analysis was performed using the `go_enrich_MF` and `go_enrich_BP` functions of clusterprofiler ([51]). The ten most represented GO categories for each analysis and their computed values can be in the All_GO-terms.pdf file found at https://osf.io/gkh89/

### MutistageOT

MutistageOT was used as per: https://github.com/dahlinlab/MultistageOT/blob/main/scripts/notebooks/real_data/paul2015/main.ipynb. We first picked the coordinates of a representative “initial cell state locus” on the UMAP using n0=3 (Supplementary figure 4B). We then defined two terminal states (Supplementary figure 4C), and applied MutistageOT to compute a pseudo-temporal order (Supplementary figure 4D) using default parameter values (n_groups = 20, epsilon = 0.055 patience = 1000 and tolerance = 1e-4). Cell types were defined as: Stem cells= cluster 6; Diff_cells_1= cluster 1; Diff_cells_2= cluster 0; Transit cells= all other clusters; for cluster definition, see Figure 1A.

### Motifs analysis

In order to analyze the over-representation or under-representation of certain motifs, we computed an empirical pvalue by permutation (see [96]). For this, we first computed B sign-randomized graphs.

When the observed number of motifs *X_obs_* was superior to the mean number of corresponding randomized motifs 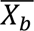, we assessed its over-representation by counting the number of times the randomized value *X_b_* was equal to or larger that the observed number of motifs *X_obs_* such that:

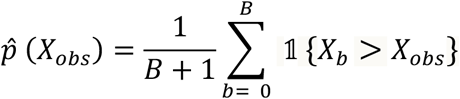

When the observed number of motifs *X_obs_* was inferior to the mean number of corresponding randomized motifs 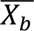, we assessed its under-representation with:

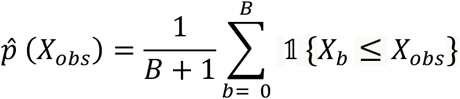

The precision of the pvalue being equal to 1/√*B*, we display the results with the according precision.

